# MEST induces epithelial-mesenchymal transition through IL-6/JAK/STAT3 signaling in breast cancers

**DOI:** 10.1101/295832

**Authors:** Min Soo Kim, Hyun Sook Lee, Yun Jae Kim, Sung Gyun Kang, Do Yup Lee, Wook Jin

## Abstract

The loss of imprinting of MEST has been linked to certain types of cancer by promoter switching. However, MEST-mediated regulation of tumorigenicity and metastasis are yet to be understood. Herein, we reported that MEST is a key regulator of IL-6/JAK/STAT3/Twist-1 signal pathway-mediated tumor metastasis. Enhanced MEST expression is significantly associated with pathogenesis of breast cancer patients. Also, MEST induces metastatic potential of breast cancer through induction of the EMT-TFs-mediated EMT program. Moreover, MEST leads to Twist-1 induction by STAT3 activation and subsequently enables the induction of activation of the EMT program via the induction of STAT3 nuclear translocation. Furthermore, the c-terminal region of MEST was essential for STAT3 activation via the induction of JAK2/STAT3 complex formation. Finally, MEST significantly increases the breast cancer’s ability to metastasize from the mammary gland to the lung. These observations suggest that MEST is a promising target for intervention to prevent tumor metastasis.

## Introduction

Mesoderm specific transcript (MEST) has been shown to maternally imprint on parthenogenetic embryos, and only the paternally inherited allele is preferentially expressed in the placenta of humans. Additionally, MEST expressed in extraembryonic tissues during development is paternally transmitted, and the offspring exhibits severe intrauterine growth retardation after a null allele, suggesting that MEST may play a role in development (Lefebvre et al, 1997; Mayer et al, 2000). Moreover, the MEST gene mapped to chromosome 7q32 and has homologous sequences on the short arm of human chromosomes 3 and 5 (Riesewijk et al, 1997). Furthermore, MEST is thought to encode a protein with homology to the alpha/beta-hydrolase superfamily, but the precise function has not been identified (Kosaki et al, 2000; Riesewijk et al, 1997).

Recently, several studies have described the frequent loss of imprinting of MEST, which has been linked to certain types of cancer. In choriocarcinoma, trophoblast tumors, breast cancer, lung adenocarcinomas, uterine leiomyoma, colon cancer, human adult blood lymphocytes, and lymphoblastoid cell lines, MEST is expressed from both the paternal allele and the maternal allele via promoter switching from isoform 1 to isoform 2 of MEST (Kosaki et al, 2000; Mayer et al, 2000; Moon et al, 2010; Nakanishi et al, 2004; Nishihara et al, 2000; Pedersen et al, 1999). Although MEST is expressed in many tumors by promoter switching, it is currently unclear how MEST regulates tumorigenicity and metastasis of the tumor. Moreover, the signaling mechanism that induces and maintains tumorigenicity and tumor metastasis by MEST has remained unclear.

In this report, we first identified a new network involved in tumor metastasis and EMT that regulates and coordinates MEST. We found that a high level of MEST expression is significantly correlated with pathogenesis in breast cancer patients and that MEST acts as a key regulator of tumor metastasis through IL-6/JAK/STAT3/Twist-1 signal pathway-mediated EMT induction and may regulate the maintenance of the CSC state.

## Results

### Elevated MEST expression in human breast tumors and cells

Although there have been previous studies showing that the loss of MEST imprinting is linked to certain types of cancer by promotor switching, the expression patterns of MEST have not been well characterized in human breast cancer tissues. To evaluate its potential involvement and role in breast cancer, we first examined human tumor specimens to determine whether MEST expression was associated with certain pathological phenotypes in clinical breast tumor samples. Strikingly, the level of expression of MEST determined by quantitative RT-PCR was significantly enhanced in 16 out of 17 tumor tissues of breast cancer patients (94%) compared to those in corresponding patient-matched normal tissue samples (**Figure 1A**). Based on these observations, we next examined MEST expression in a panel of established tumorigenic or metastatic mouse and human breast cancer cells. MEST was highly expressed in tumorigenic and metastatic human breast cancer cells compared to the level in human mammary epithelial cells (HMLE) and nontumorigenic MCF10A cells. Nine of the eleven metastatic breast cancer cells expressed MEST protein, whereas only two of the four non-metastatic breast cancer cells expressed MEST protein. Additionally, we found that MEST expression was up-regulated in 4T1 cells, the most aggressive of the three mouse mammary tumor cells and the only one capable of forming macroscopic lung metastases (**Figures 1B** and **S1**). These observations suggested that MEST is often induced during tumor progression. Next, MEST expression was evaluated by immunohistochemistry analysis in a series of 59 human tumor tissues of breast cancer patients, and a significant correlation was found between MEST expression and pathological phenotypes. Normal breast tissue samples exhibited low expression of MEST. However, elevated MEST expression was detected in the infiltrating ductal (IDC) and metastatic breast carcinoma samples (**Figure 1C**).

**Figure 1.**
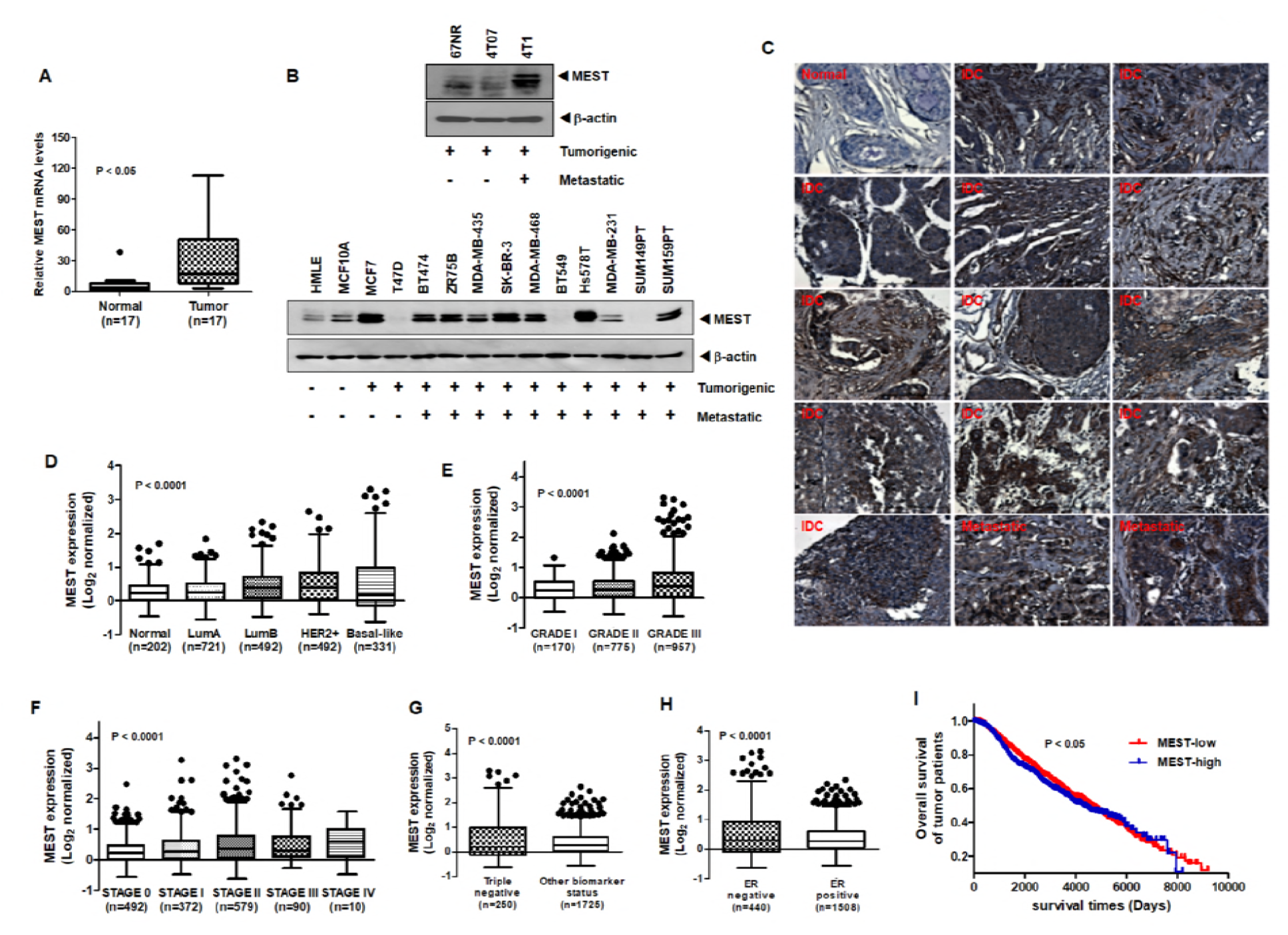
Expression of MEST is correlated with breast pathological phenotypes. (A) The expression of the mRNA encoding MEST in 17 samples of individual human breast cancer patient relative to that of the healthy tissue, as determined by quantitative RT-PCR. The 18S mRNA level was used to normalize the variability in template loading. *P* < 0.05, *t*-test. (B) Western blot analysis of MEST expression in tumorigenic and metastatic human or mouse breast cancer cells. β-actin was used as a loading control. (C) Representative immunohistochemical images of MEST staining in normal human breast tissue, infiltrating duct carcinoma (IDC), and metastatic breast carcinoma (magnification: 200×). (D) Box-and-whisker (Tukey) plots are shown for the expression of MEST in 2,137 human breast cancer patients by subtypes of breast cancer. The MEST level was extracted from the METABRIC dataset and averaged for each tumor. *P* < 0.0001, ANOVA. (E) Box-and-whisker (Tukey) plots are shown for the expression of MEST on the grade of 2,137 human breast cancer patients. (F) Box-and-whisker (Tukey) plots are shown for the expression of MEST on the stage of 2,137 human breast cancer patients. *P* < 0.0001, ANOVA. (G) Box-and-whisker (Tukey) plots are shown for the expression of MEST on triple negative or other biomarkers from 2,137 human breast cancer patients. *P* < 0.0001, *t*-test. (H) Box-and-whisker (Tukey) plots are shown for the expression of MEST on ERα negative or ERα positive samples from 2,137 human breast cancer patients. The MEST level from Figure 1D, 1E, 1F, 1G, and 1H was extracted from the METABRIC dataset and averaged for each tumor. Points below and above the whiskers are drawn as individual dots. *P* < 0.05, *t*-test. (I) Kaplan-Meier survival plot of low and high MEST expression on the METABRIC data set. Patients were divided into high- and low-MEST expressers, and their survivals were compared. The *P* value was calculated by a log-rank test (*P* < 0.05).

To further examine the relationship between MEST expression and breast cancer progression, we investigated the clinical relevance using MEST publicly available gene expression microarray datasets of breast cancer patients from 2,137 breast cancer patients in the METABRIC dataset and 855 breast cancer patients from the UNC dataset (GSE26338) (Curtis et al, 2012; Harrell et al, 2012). Consistent with the above observations, MEST expression was significantly higher in all breast cancer subtypes compared to normal breast tissues and strongly correlated with breast cancer subtypes (**Figures 1D** and **S2A**). Specifically, the level of MEST expression was significantly higher in both basal-like and claudin-low breast cancer subtypes, and moreover, MEST expression was correlated with the grade and stage of breast cancer patients (**Figures 1E**, **1F**, **S2B**, and **2C**). Furthermore, MEST expression was significantly upregulated in triple-negative breast cancer (TNBC) relative to the luminal and HER2-positive markers in these independent datasets (**Figures 1G** and **S2D**), and in particular, MEST expression was strongly correlated with ER-α expression (**Figures 1H** and **S1E**). This finding is consistent with features of the basal-like and claudin-low breast cancer subtypes, which are known to be closely correlated with the subset of receptor-negative breast cancers.

In a recent discovery, basal-like and claudin-low breast cancer subtypes have been linked with resistance to chemotherapy, acquisition of stem cell characteristics, metastasis, and poor survival (Hennessy et al, 2009). For these reasons, we performed clinical prediction analyses on the breast tumors in the METABRIC dataset and UNC dataset. As a result, it seems that a high level of MEST expression could be a good predictor of poor clinical outcome. Those patients whose tumors exhibited high levels of MEST expression showed a significantly poorer survival outcome than those with tumors that exhibited low levels of MEST expression, which correlated with the metastatic free survival status in the UNC dataset (**Figures 1I** and **S2F**). Our observations indicated that MEST might play an important role in the metastatic dissemination of breast cancer.

### MEST enhances tumorigenicity and metastatic potential of breast cancer

The invasion-metastasis cascade is driven by the acquisition of anchorage independence, gain of motility, invasion in and out of circulation, and colonization of distant organs (Fidler, 2003; Hanahan & Weinberg, 2011; Valastyan & Weinberg, 2011). Based on the above observation, we speculated that MEST might contribute to tumorigenicity and the metastasis process of breast cancer. To test this notion, we first selected highly metastatic Hs578T, SUM159PT, and 4T1 cells and engineered stably expressing MEST-shRNAs into the cell lines. As shown in Figure 2A, MEST-shRNA suppressed the expression of endogenous MEST by 80% (**Figure 2A**).

**Figure 2.**
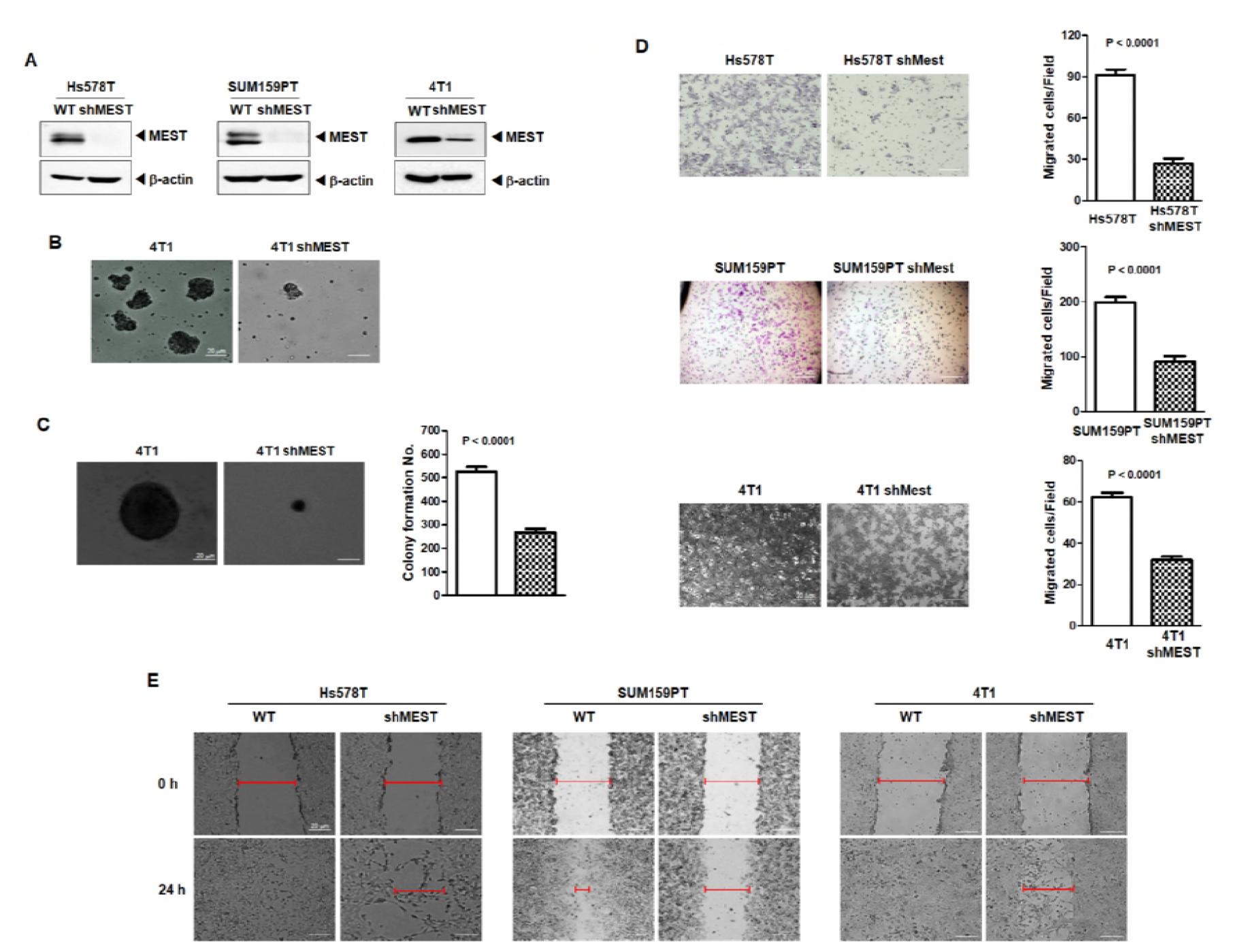
MEST induces tumorigenic and metastatic potential of breast cancer. (A) Western blot analysis of MEST expression in Hs578T, SUM159PT, and 4T1 control-shRNA or MEST-shRNA cells. β-actin was used as a loading control. (B) Formation of spheroid colonies of 4T1 control-shRNA or MEST-shRNA cells. Spheroid colonies in suspension were then photographed at 200× magnification. (C) The soft agar colony-forming assay of the 4T1 control-shRNA or MEST-shRNA cells (n=3). *P* < 0.0001, *t*-test. (D) The migration assay of in Hs578T, SUM159PT, and 4T1 control-shRNA or MEST-shRNA cells (n=3). *P* < 0.0001, *t*-test. (E) Wound healing assay of Hs578T, SUM159PT, and 4T1 control-shRNA or MEST-shRNA cells. Wound closures were photographed at 0 and 24 hrs after wounding.

By an anoikis assay measuring the anchorage-independent survival (Jin et al, 2010), we examined the clonal formation abilities of 4T1 control-shRNA and 4T1 MEST-shRNA cells in suspension. To accomplish this, we used cell culture dishes to determine if 4T1 control-shRNA and 4T1 MEST-shRNA cells could not attach. Under these conditions, 4T1 control-shRNA cells proliferated as large spheroid aggregates in suspension, whereas 4T1 MEST-shRNA cells exhibited a significantly lower survival outcome in suspension (**Figure 2B**).

We next examined whether the loss of MEST affected the colony forming ability of 4T1 cells. 4T1 control-shRNA cells exhibited almost 2-fold higher colony-forming activity compared to the 4T1 MEST-shRNA cells (**Figure 2C**). To test for other functional hallmarks of metastatic potential and mesenchymal/CSC state, we conducted cell motility and wound healing assays. Hs578T, SUM159PT, 4T1 control-shRNA, and MCF10A-MEST overexpressing cells had significantly increased cell migration, whereas MEST-shRNA cells and MCF10A control cells inhibited the migration of breast cancer cells, indicating that the increased cell migration was correlated with the level of MEST expression (**Figures 2D, 2E** and **S3**). These findings suggest that MEST accelerates the secondary tumor forming ability to distant organ sites.

### MEST regulates the invasion-metastasis cascade through induction of a Twist-mediated EMT program

Since the epithelial-mesenchymal transition (EMT) broadly regulates invasion and metastasis by acquiring cellular traits associated with high-grade malignancy (Hanahan & Weinberg, 2011; Valastyan & Weinberg, 2011), and induction of the EMT program has been strongly associated with the acquisition of self-renewal traits (Mani et al, 2008; Scheel et al, 2011), we speculated that the acquisition of tumorigenic and metastatic potential by MEST expression might depend on induction of the EMT program. To address the issue, we examined whether the expression of MEST regulates an EMT program in breast cancer cells. Specifically, compared to the level of expression in Hs578T, SUM159PT, and 4T1 control-shRNA cells, the cell lines expressing MEST-shRNA exhibited increased expression of E-cadherin protein and significant decreases in N-cadherin, fibronectin, and vimentin expression (**Figure 3A**). Moreover, quantitative RT-PCR analyses revealed that the expression of E-cadherin in Hs578T, SUM159PT, and 4T1 control-shRNA cells was strongly decreased, compared to the level of expression in Hs578T, SUM159PT, and 4T1 MEST-shRNA cells, while the expression levels of mRNAs encoding mesenchymal markers were markedly increased, which correlated with protein levels (**Figure S4A**).

**Figure 3.**
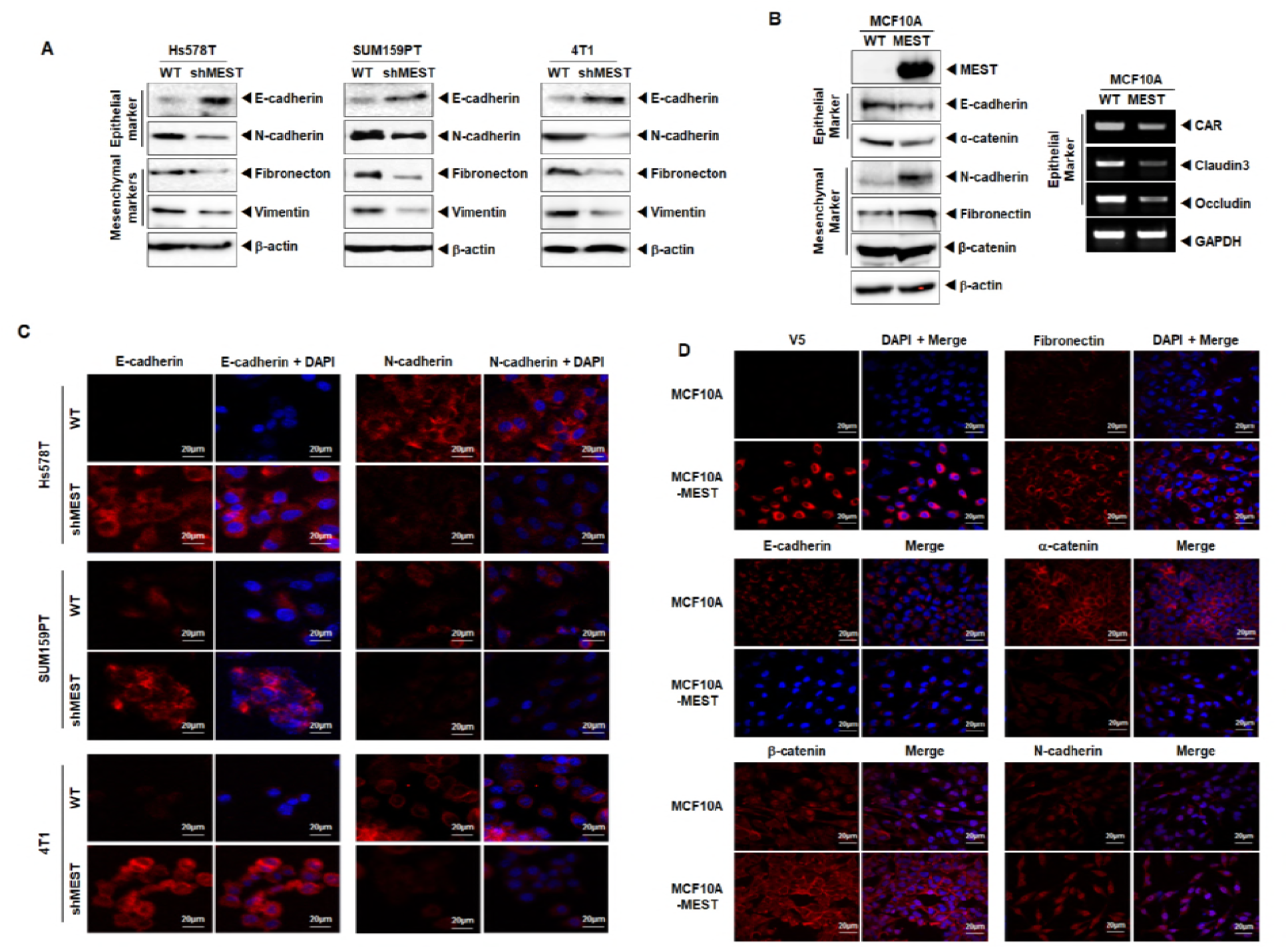
High MEST expression induces EMT program. (A) Western blot analysis of the expression of E-cadherin, N-cadherin, fibronectin, and vimentin proteins in Hs578T, SUM159PT, and 4T1 control-shRNA or MEST-shRNA cells. β-actin was used as a loading control. (B) Western blot (Left) and RT-PCR (Right) analysis of the expression of CAR, caludin3, occludin, E-cadherin, α-catenin, N-cadherin, fibronectin, and β-catenin proteins in MCF10A control or MCF10A-MEST cells. GAPDH and β-actin were used as a loading control. (C) Immunofluorescence images of E-cadherin and N-cadherin in Hs578T, SUM159PT, and 4T1 control-shRNA or MEST-shRNA cells. The red or green signal represents staining of the corresponding protein, while the blue signal represents DAPI staining. (D) Immunofluorescence images of V5, E-cadherin, β-catenin, fibronectin, α-catenin, and N-cadherin in MCF10A control or MCF10A-MEST cells. The red signal represents the staining of the corresponding protein, while the blue signal represents DAPI staining.

To further assess whether MCF10A-overexpressing MEST cells retain the gene expression profiles observed in highly metastatic breast cancer cells that passed through EMT, we examined the expression profile of multiple EMT-associated proteins. We found the expression levels of epithelial and mesenchymal markers in MCF10A-MEST cells were similar with those in Hs578T, SUM159PT, and 4T1 control-shRNA cells. Specifically, relative to levels in the MCF10A control cells, MCF10A-MEST cells exhibited a significant increase in the expression of E-cadherin, or α-catenin protein, and decreases in the expression of N-cadherin, fibronectin, β-catenin and vimentin proteins (**Figure 3B**). Moreover, RT-PCR and quantitative RT-PCR analyses revealed that the mRNA expression levels of the epithelial markers CAR, Claudin3, Occludin and E-cadherin (~50-fold) in MCF10A-MEST cells were strongly decreased, while the mRNA expression levels of mesenchymal markers were significantly increased, specifically N-cadherin (~330-fold), Vimentin (~10-fold), and Fibronectin (~116-fold) (**Figures 3B** and **S4B**).

We also performed immunostaining to further determine whether MEST expression induced the EMT program. The immunostaining intensity of epithelial or mesenchymal markers in Hs578T, SUM159PT, and 4T1 control-shRNA cells corresponded to the level of protein and mRNA expression in the cells (**Figure 3C**). Upon MEST knockdown, the expression of E-cadherin was greatly enhanced, while the expression of mesenchymal markers, such as N-cadherin, was significantly decreased, indicating that the depletion of MEST prevents the conversion of mesenchymal traits on epithelial cells. In addition, the immunostaining analysis revealed that the expression levels of E-cadherin and α-catenin in the MCF10A-MEST cells were significantly decreased, while N-cadherin, fibronectin, and β-catenin expression levels were markedly increased (**Figure 3D**), compared to the expression level in MCF10A control cells.

During the process of tumor metastasis, a set of pleiotropically acting transcription factors, including Goosecoid, Foxc1, Foxc2, Slug, SIP1, Twist-1, and Twist-2, orchestrate the EMT program and related migratory processes. Additionally, these pleiotropically acting transcription factors are expressed in various combinations in a number of malignant tumor types (Hanahan & Weinberg, 2011; Mani et al, 2008). These recent studies and the above observation led us to speculate that MEST might contribute to the induction of EMT via the upregulation of transcriptional regulators. To test this possibility, we examined the expression of a set of pleiotropically acting transcription factors. There was a considerable increase in the expression of EMT-inducing transcription factors, specifically Goosecoid, Foxc1, Foxc2, and Twist-1, in Hs578T, SUM159PT, and 4T1 control-shRNA cells but not Slug, SIP1, and Twist-2, compared to the expression in Hs578T, SUM159PT, and 4T1 MEST-shRNA cells (**Figure 4A**). The difference was also confirmed by quantitative RT-PCR. MCF10A-MEST cells overexpressed the EMT-inducing transcription factors, such as Foxc1 (16-fold), Foxc2 (50-fold), and Twist-1 (15-fold), but not Slug, SIP1, and Twist-2 (**Figure S5A**), compared to the MCF10A control cells. These data indicate that the overexpression of MEST upregulates the expression of multiple genes in EMT-induced cells and MEST may be a key mediator of *twist* gene trans-activation, resulting in the loss of epithelial characteristics and the gain of mesenchymal properties.

**Figure 4.**
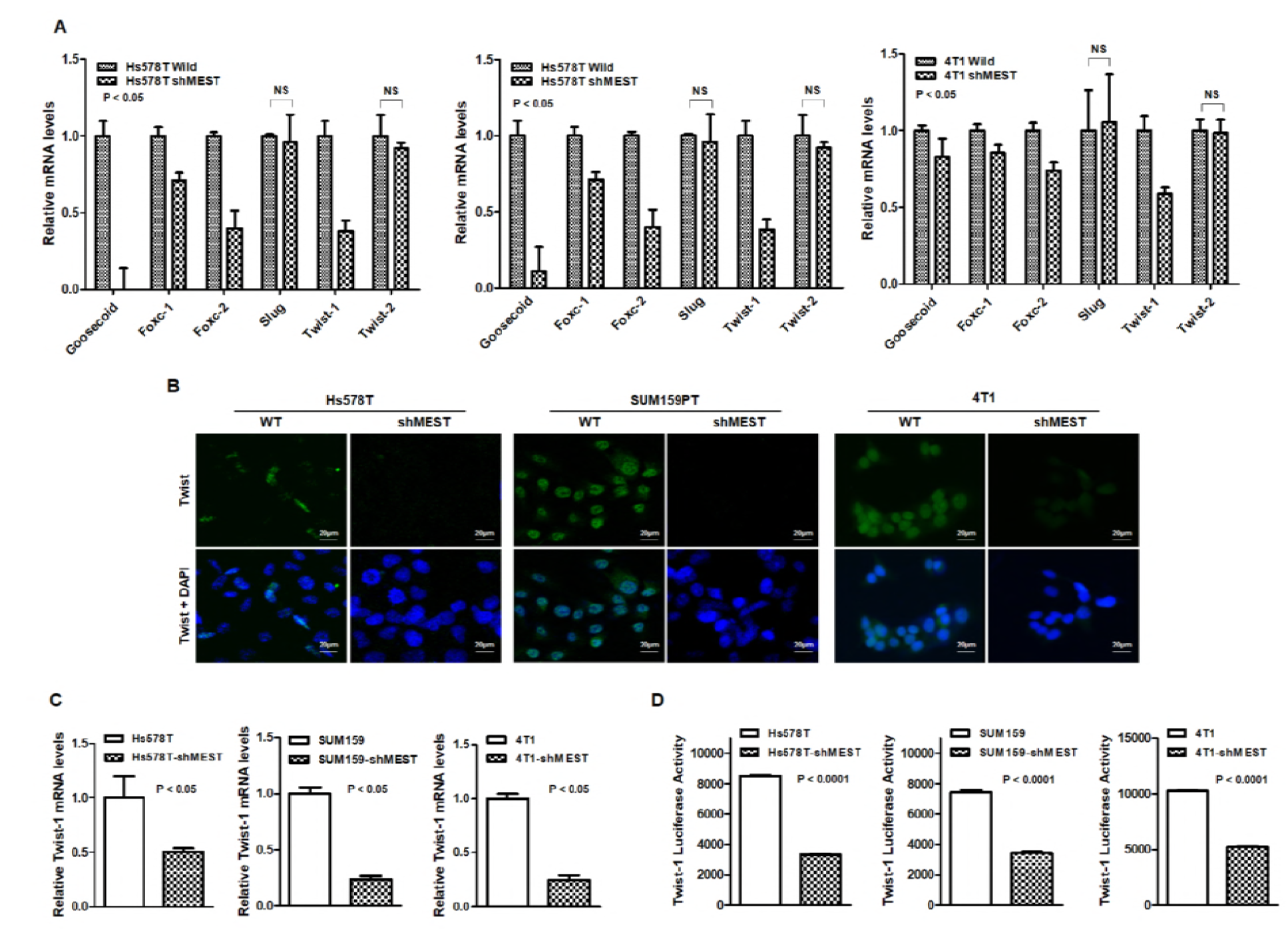
High MEST expression induces EMT through upregulation of EMT-inducing transcription factors. (A) The relative expression levels of mRNA encoding Goosecoid, Foxc-1, Foxc-2, Slug, SIP-1, Twist-1, and Twist-2 in Hs578T, SUM159PT, and 4T1 control-shRNA or MEST-shRNA cells, as determined by quantitative RT-PCR. 18S was used as a loading control. *P* < 0.05, *t*-test. (B) Immunofluorescence images of Twist-1 in Hs578T, SUM159PT, and 4T1 control-shRNA or MEST-shRNA cells. The green signal represents the staining of the corresponding protein, while the blue signal represents DAPI staining. (C) The relative expression levels of mRNA encoding Twist-1 in Hs578T, SUM159PT, and 4T1 control-shRNA or MEST-shRNA cells, as determined by quantitative RT-PCR. 18S was used as a loading control. *P* < 0.05, *t*-test. (D) Promoter activity of Twist-1 gene in Hs578T, SUM159PT, and 4T1 control-shRNA or MEST-shRNA cells were measured by the Luciferase reporter assay. Each bar represents the mean ± SEM of three experiments. *P* < 0.0001, *t*-test.

Because MEST more significantly regulates Twist-1 expression compared to other EMT-inducing transcription factors (**Figure 4A**), we initially focused on the upregulation of Twist-1 by MEST expression. Immunofluorescence studies confirmed that Twist in Hs578T, SUM159PT, 4T1 control-shRNA cells, and MCF10A-MEST cells was expressed primarily as a nuclear form of Twist (**Figures 4B** and **S5B**). The intensity of overlap between Twist immunofluorescence and DAPI significantly decreased in MEST knockdown cells and MCF10A control cells, compared to those in Hs578T, SUM159PT, 4T1 control-shRNA cells, and MCF10A-MEST cells. To test the possibility of the role of MEST in Twist regulation at the transcriptional level, a quantitative RT-PCR analysis and Twist promoter activity assay were carried out in the control and MEST knockdown cells. Accumulation of *twist* mRNA was diminished in MEST knockdown cells (**Figure 4C**) and a significant reduction of *Twist* promoter activity was also observed in MEST knockdown cells (**Figure 4D**). Collectively, these data imply that MEST is a positive mediator of *twist* gene trans-activation, resulting in the loss of epithelial characteristics and the gain of mesenchymal properties.

### MEST upregulates Twist-1 expression through activation of STAT3

Although gene ontology (GO) analysis from the UniProtKB/Swiss-Prot database proposed that MEST localizes to the endoplasmic reticulum (ER), MEST has not been shown to localize to or be associated with organelles and specialized subcellular compartments. Since subcellular localization is important for functionality, we examined the subcellular localization of MEST. It was found that the majority of MEST protein was present in the membrane fraction, including the plasma/ER/Golgi/mitochondrial membranes, and a small fraction of MEST protein was found in the nucleus where the Twist protein was primarily located. It is worth noting that cytokeratin 18 expression is well-known as a luminal epithelial marker and was markedly increased in the SUM159PT-shMEST cells compared to the control-shRNA cells (**Figures S6A** and **S6B**). This result supports that MEST regulates the invasion-metastasis cascade through induction of the Twist-mediated EMT program. However, the distinction in the subcellular localization of MEST and Twist led us to hypothesize that MEST might play a role as a linker or scaffold protein having characteristics of both nuclear and cytoplasmic signaling molecules. Recently, Cheng *et al*. (Cheng et al, 2008) demonstrated that the active form of STAT3 was able to directly bind to the *Twist* promoter and promote its transcriptional activity. These observations led us to speculate that MEST might be involved in the regulation of STAT3 activation and that it was functionally linked to the regulation of Twist-1. To test the effect of MEST in STAT3 activation, we initially examined whether knockdown of MEST in the Hs578T, SUM159PT, and 4T1 cells affected both the total and active forms of STAT3 protein. We found that STAT3 activation, as well as Twist-1 expression, was markedly decreased in the Hs578T, SUM159PT, and 4T1-shMEST cells relative to its control-shRNA cells; however, STAT3 expression was not affected in the knockdown of MEST. Moreover, the levels of phosphorylated and total Jak2 were not altered upon MEST knockdown (**Figure 5A**). In addition, similar results were obtained with MCF10A-MEST cells. STAT3 phosphorylation and Twist-1 expression were significantly increased, but JAK2 phosphorylation, total JAK2, and STAT3 expression were not changed (**Figure S6C**).

**Figure 5.**
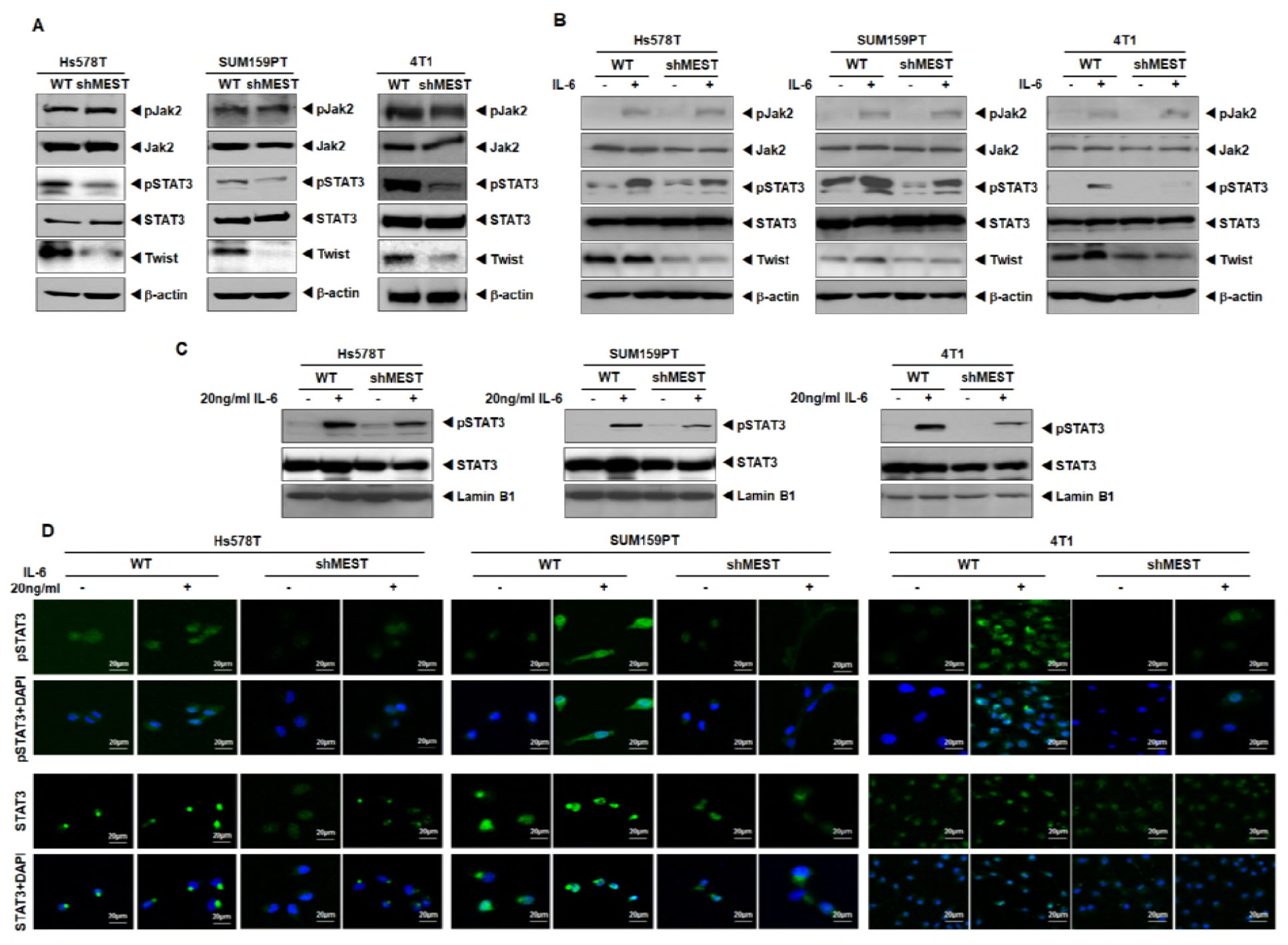
MEST led to Twist-1 upregulation through activation of STAT3. (A) Western blot analysis of the expression of phospho-JAK2, JAK2, phospho-STAT3, STAT3, and Twist-1 proteins in Hs578T, SUM159PT, and 4T1 control-shRNA or MEST-shRNA cells. β-actin was used as a loading control. (B) Western blot analysis of the expression of phospho-JAK2, JAK2, phospho-STAT3, STAT3, and Twist-1 proteins in Hs578T, SUM159PT, and 4T1 control-shRNA or MEST-shRNA cells treated with or without 20 ng/ml IL-6. β-actin was used as a loading control. (C) Western blot analysis of the expression of phospho-STAT3 and STAT3 proteins in Hs578T, SUM159PT, and 4T1 control-shRNA or MEST-shRNA cells after nuclear fractionation. Lamin B1 was used as a nuclear loading control. (D) Immunofluorescence images of phospho-STAT3 and STAT3 in Hs578T, SUM159PT, and 4T1 control-shRNA or MEST-shRNA cells. The green signal represents the staining of the corresponding protein, while the blue signal represents DAPI staining.

Since Jak2 expression and activation were similar between control and MEST knockdown cells, we examined whether there was any difference in terms of ligand-induced Jak2-STAT3 activation between the control and MEST knockdown cells or the levels of Twist were affected by ligand-induced STAT3 activation in MEST knockdown cells. To test this notion, we assessed the expression level and activation of JAK2 of Hs578T, SUM159PT, and 4T1control-shRNA and MEST-shRNA cells with IL-6 activation. Interestingly, there was no difference in the expression or activation of JAK2 between the control and MEST knockdown cell and no difference in STAT3 expression after stimulation with or without IL-6. However, IL-6-induced STAT3 activation was markedly reduced in MEST knockdown cells, relative to the control shRNA cells, which was strongly correlated with the level of Twist-1 expression (**Figure 5B**).

Since activated STAT3 can translocate into the nucleus and act as a transcription factor, we consequently tested whether MEST could modulate IL-6-dependent subcellular localization of STAT3. In Hs578T, SUM159PT, and 4T1control-shRNA cells, we observed an increase in the nuclear translocation and nuclear expression of STAT3 and phospho-STAT3 with or without IL-6 treatment, compared to those in MEST-shRNA cells (**Figures 5C** and **5D**). These results suggest that treatment of IL-6 could induce STAT3 phosphorylation in control cells; however, the loss of MEST seemed to have inhibitory effects on STAT3 phosphorylation after IL-6 treatment. Taken together, these observations indicate that MEST regulates expression of Twist, an EMT transcription factor, possibly mediated through JAK2 activation-independent STAT3 activation.

Although we did not find a decrease in total STAT3 levels in MEST knockdown cells, we sought to test the possibility of a regulatory role of MEST in STAT3 activation at the transcriptional level. The *STAT3* mRNA expression and *STAT3* promoter activity were not affected by IL-6 treatment in both the control and MEST knockdown cells (**Figures S7A** and **S7B**), indicating that MEST-mediated STAT3 activation is not transcriptionally regulated.

### MEST induces STAT3 activation through MEST-STAT3 complex formation

To test the regulatory role of MEST in STAT3 activation, we attempted to determine whether MEST regulates STAT3 activation through MEST-STAT3 complex formation by investigating the complex formation of MEST and STAT3 endogenously. An interaction between these two proteins was readily detected. Reciprocal immunoprecipitation using STAT3 antibody confirmed the presence of the MEST-STAT3 complex at the endogenous level (**Figure 6A**). Moreover, MEST directly interacts with STAT3 under physiological conditions (**Figure 6B**). To identify the STAT3 functional domain responsible for its interaction with MEST, we used a series of deletion constructs of STAT3. A STAT3 mutant containing the N-terminal domain (NTD: protein interaction domain) interacted with MEST, whereas MEST did not interact with SH2 or the transactivation domain of STAT3 (**Figures 6C** and **S8A**). MEST protein is annotated as a putative multi-pass membrane protein by sequence prediction (**Figure 6D**). To identify the MEST functional regions responsible for its interaction with STAT3, we used a deletion construct of MESTΔC and full-length MEST. Full-length MEST interacts with STAT3, whereas MESTΔC with a C-terminal deletion after the 287^th^ amino acid did not interact with STAT3 (**Figure 6D**).

**Figure 6.**
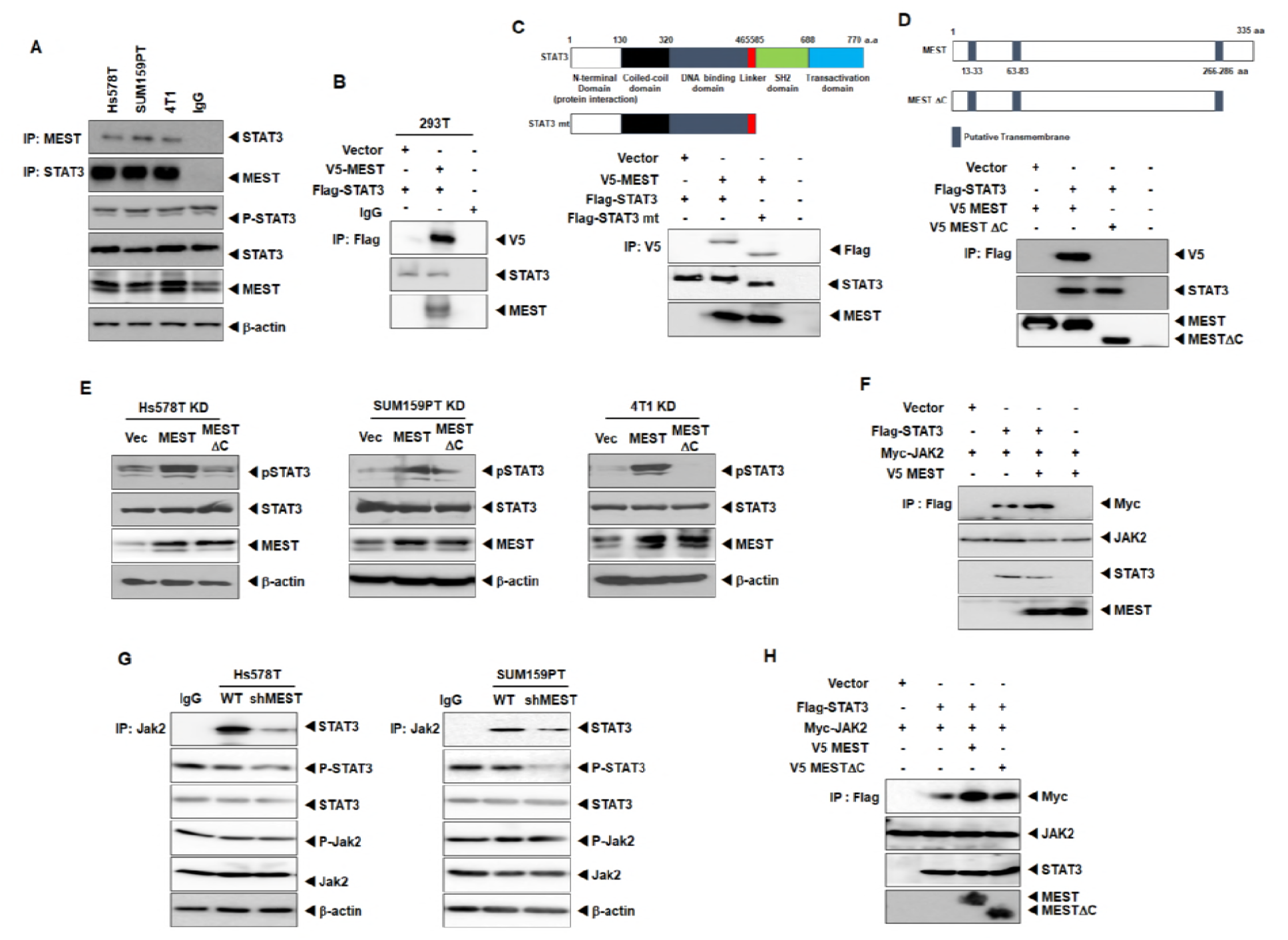
MEST induces STAT3 activation through induction of JAK2-STAT3 complex formation. (A) Identification of the complex formation of endogenous MEST/STAT3 in MBA-MB-231 and Hs578T cells. (B) Western blot analysis of the whole-cell lysates and immunoprecipitates derived from HEK293 cells transfected with the V5-MEST and Flag-STAT3 constructs, as indicated. (C) Identification of the STAT3 region that interacted with MEST. Western blot analysis of whole-cell lysates and immunoprecipitates derived from 293T cells transfected with V5-MEST, Flag-STAT3 wild and the STAT3 deletion construct (STAT3 mt) as indicated. (D) Western blot analysis of the whole-cell lysates and immunoprecipitates derived from 293T cells transfected with the V5-MEST, V5-MESTΔC, and Flag-STAT3 constructs, as indicated. (E) Western blot analysis of the expression of phospho-STAT3 and STAT3 proteins in Hs578T, SUM159PT, and 4T1 MEST-shRNA cells after transient transfection of MEST or MESTΔC constructs. β-actin was used as a loading control. (F) Western blot analysis of whole-cell lysates and immunoprecipitates derived from 293T cells after transfection of JAK2, STAT3, or MEST constructs. β-actin was used as a loading control. (G) Identification of the complex formation of endogenous JAK2/STAT3 in HS578T and Hs578T, SUM159PT cells or MEST-shRNA cells. (H) Western blot analysis of whole-cell lysates and immunoprecipitates derived from 293T cells after transfection of JAK2, STAT3, MEST or MESTΔC constructs. β-actin was used as a loading control.

Based on the above observation, we speculated that MESTΔC might contribute to activation of STAT3. To test this notion, we assessed STAT3 expression and STAT3 activation in MEST knockdown cells after transient transfection of MEST or MESTΔC. Compared to those in the MEST knockdown cells expressing the control or MESTΔC, the introduction of MEST to the MEST knockdown cells showed significantly increased activation and expression of STAT3 or Twist-1 (**Figures 6E** and **S8B**). These results indicated that the C-terminal region (amino acid 287-335) of MEST is essential for STAT3 activation.

Next, we attempted to determine whether MEST regulates JAK2/STAT3 complex formation through the interaction of MEST and STAT3. Interestingly, MEST induces JAK2/STAT3 complex formation (**Figures 6F** and **6G**). To identify the MEST functional region responsible for the induction of JAK2/STAT3 complex formation, we tested JAK2/STAT3 complex formation by MEST and MESTΔC. JAK2/STAT3 complex formation by MESTΔC still remained, but the level of JAK2/STAT3 interaction was significantly decreased relative that of MEST (**Figure 6H**). These findings suggest that MEST is essential for the induction of JAK2/STAT3 complex formation.

### MEST expression significantly increases metastasis in vivo

To determine whether MEST knockdown affected the ability of 4T1 cells to metastasize, 4T1 control-shRNA or MEST-shRNA cells were injected into the tail vein of BALB/c mice, and their lungs were examined for metastases 29 days after injection. The 4T1 MEST-shRNA cells strongly reduced the number of metastatic nodules relative to the 4T1 control-shRNA cells (**Figure 7A**). Additionally, a few metastatic nodules in the lungs of mice carrying MEST-shRNA cells retained MEST expression, but MEST expression in the lungs of mice with MEST-shRNA cells was greatly reduced relative to the lungs of mice carrying control-shRNA cells (**Figures 7B** and **7D**). Moreover, histological analyses confirmed that the number of micrometastatic lesions was drastically reduced in the lungs of mice with 4T1 MEST-shRNA cells (**Figure 7C**).

**Figure 7.**
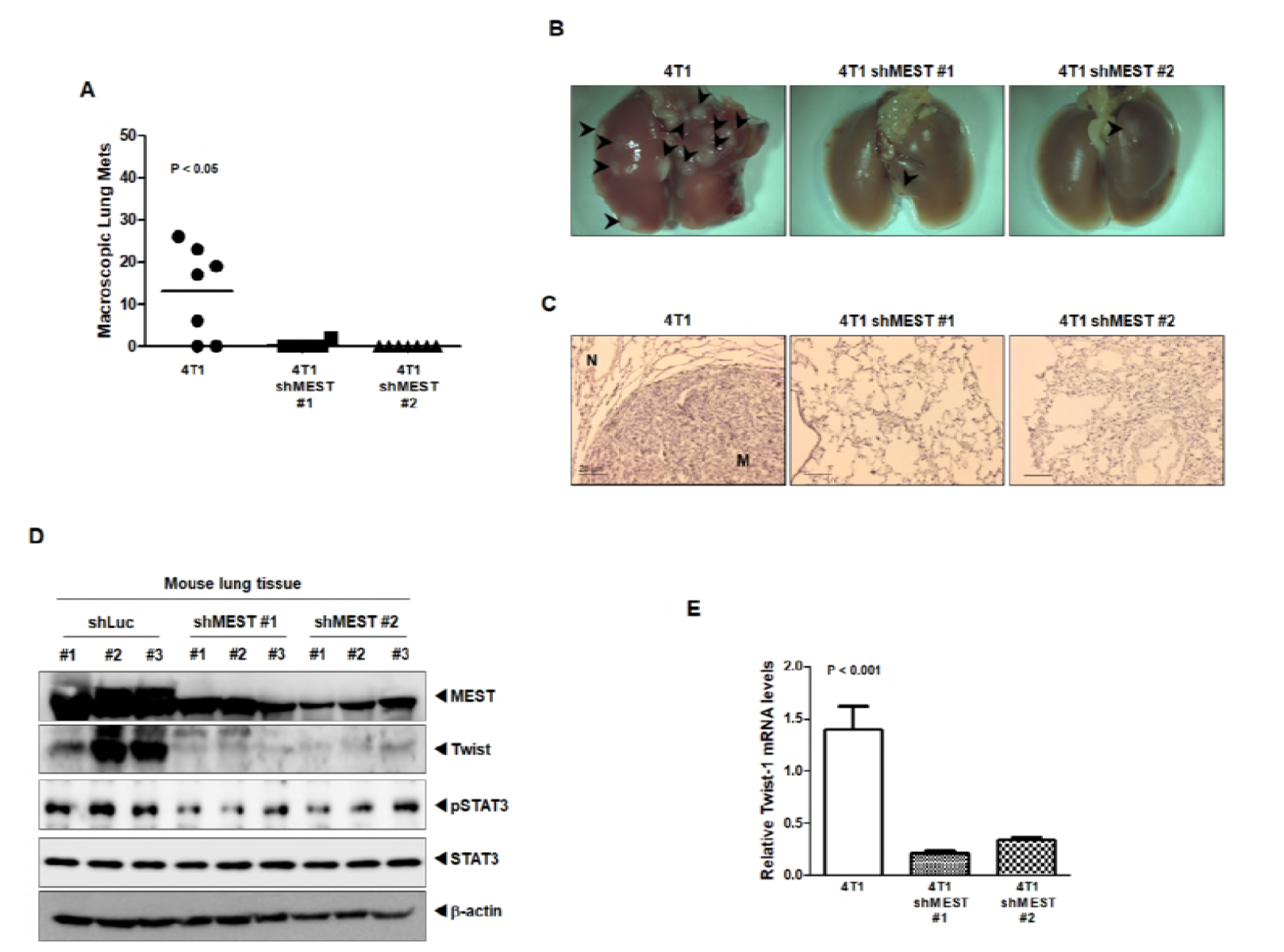
Suppression of MEST inhibited the ability of 4T1 cells to metastasize to the lung. (A) Lung metastasis by 4T1 control-shRNA or 4T1 MEST-shRNA cells. The total numbers of lung metastatic nodules from each mouse harboring 4T1 tumors expressing control-shRNA or MEST-shRNA were counted using a dissection scope (n = 6 mice/group). *P* < 0.05, ANOVA. (B) Representative images of the lungs from the mice harboring 4T1 control-shRNA or 4T1 MEST-shRNA cells for 30 days after implantation of mouse mammary fat pads of mice. (C) Representative H&E staining in the sections of the lungs from Figure 7B. N, Normal lung tissue; M, metastatic nodule. (D) Western blot analysis of MEST, phospho-STAT3, STAT3, and Twist-1 in tumor cells recovered from the lungs of individual mice expressing either 4T1 control-shRNA or 4T1 MEST-shRNA. β-actin was used as a loading control. (E) Quantitative RT-PCR of Twist-1 in tumor cells recovered from the lungs of individual mice expressing either 4T1 control-shRNA or 4T1 MEST-shRNA. 18S mRNA was used as a loading control. *P* < 0.001, *t*-test.

We speculated that the presence of small numbers of nodules in the lungs of mice carrying 4T1 MEST-shRNA cells was caused by incomplete knockdown of MEST. To test this notion, we examined the level of MEST expression in the lungs of mice carrying 4T1 control-shRNA or MEST-shRNA cells. MEST expression in the lungs of mice with 4T1 MEST-shRNA cells was significantly lower than that of the 4T1 control-shRNA cells. Moreover, STAT3 activation and Twist-1 expression was markedly reduced in the lungs of mice with 4T1 MEST-shRNA cells relative to that of 4T1 control-shRNA cells (**Figure 7D**). Furthermore, quantitative RT-PCR analysis of Twist-1 revealed reduced expression in the lungs of mice injected with 4T1 MEST-shRNA cells compared to their control counterparts (**Figure 7E**). These results indicated that expression of MEST is required for the full metastatic ability of highly metastatic 4T1 breast cancer cells.

## Discussion

Although MEST expression has been identified in breast cancer by its promoter switching, it is unclear whether MEST is oncogenic and what is the function of MEST is in the tumorigenicity and metastasis of breast cancer. In this study, we found that MEST is frequently overexpressed in highly metastatic human and mouse breast cancer cells and clinical breast tumor samples relative to normal epithelial cells and tissue. Moreover, MEST expression was significantly associated with the subtypes, grade, stage, and triple-negative phenotype of breast cancer patients. Our observations are consistent with other previous studies, which showed that Slug, HOXB13, HER2/neu, TrkA, TrkC, Foxc2, and Goosecoid, which promote metastatic behaviors and poor prognosis in breast cancer, are already overexpressed in breast cancer patients and breast cancer cells before the appearance of the malignant tumor phenotype (Hartwell et al, 2006; Kim et al, 2013a; Kim et al, 2013b; Ma et al, 2003; Mani et al, 2007). In addition, our results suggest that MEST expression may correlate with the characteristics of metastatic breast cancer.

Metastatic breast cancer is an aggressive, chemoresistant tumor and forms a distinct subtype that is closely related to a receptor-negative breast cancer, such as basal and claudin-low cells characterized by increased motility, poor survival outcome, and the resistance of apoptosis (Hennessy et al, 2009). In our results, suppression of MEST in highly metastatic breast cancer cells results in increased anoikis and the loss of motility and invasion potential. These observations indicate that high MEST expression enables breast cancer to acquire cellular traits associated with high-grade malignancy.

Recent studies demonstrated that metastatic breast cancer is closely linked to the EMT program. Metastatic breast cancer has an enriched EMT core signature that is associated with invasion, migration, resistance to chemotherapy, as well as the acquisition of mesenchymal and cancer stem cell (CSC) traits. In response to the EMT program, EMT-inducing transcription factors are induced that orchestrate the EMT program (Hanahan & Weinberg, 2011; Mani et al, 2008; Scheel et al, 2011; Taube et al, 2010). In our present work, high MEST expression induces the EMT program. MEST expression significantly reduced the expression of epithelial markers (E-cadherin, α-catenin, CAR, claudin3, and occludin) and increased the expression of mesenchymal markers (N-cadherin, fibronectin, β-catenin and vimentin). Moreover, MEST expression increased the expression levels of EMT-inducing transcription factors, such as Goosecoid, Foxc1, Foxc2, and Twist-1, which correlates strongly with the poor survival of breast cancer. Our observations suggest that MEST expression is critical to EMT induction through EMT-inducing transcription factors and governs the conversion to a high degree of plasticity.

The IL-6/JAK2/STAT3 pathway induces mesenchymal/CSC traits via the induction of Twist-1, an EMT-inducing transcription factor, and resistance to chemotherapy (Creighton et al, 2009; Frank et al, 2010; Li et al, 2008; Liu & Wicha, 2010; Marotta et al, 2011; Rosen & Jordan, 2009). The molecular mechanisms of MEST-mediated IL-6/JAK2/STAT3 modulation in breast cancer are unclear, and none of the findings reported to date hinted at a link between MEST expression and IL-6/JAK2/STAT3 modulation. We found that upregulation of Twist-1 expression and the activation of the IL-6/JAK2/STAT3 pathway through MEST-STAT3 complex formation induced tumorigenicity and metastatic potential of breast cancer.

STAT3 activation by IL-6 leads to CSC self-renewal traits, induction of EMT, and cell transformation of a number of oncogenes (Buettner et al, 2002; Haura et al, 2005; Levy & Darnell, 2002). Additionally, Twist-1 is the master regulator of embryonic morphogenesis, playing a key role in metastasis, acquisition of CSC traits and the induction of EMT in breast cancer (Ansieau et al, 2008; Kwok et al, 2005; Ohuchida et al, 2007; Raval et al, 2005; Valsesia-Wittmann et al, 2004; Yang et al, 2004; Zhang et al, 2008; Zhang et al, 2007). Our studies further uncovered MEST function as a key regulator of Twist-1 expression via the activation of IL-6/JAK2/STAT3 signaling. In the presence of IL-6, the induction of STAT3 nuclear translocation by MEST promotes EMT through the increased expression of Twist-1.

In previous studies, MEST has been shown to possess a putative alpha/beta-hydrolase domain (Kosaki et al, 2000; Riesewijk et al, 1997). However, the function of MEST has not been adequately explored. Additionally, MEST protein may have a putative multi-pass membrane protein according to gene ontology (GO). In our hands, the MESTΔC deletion mutant containing amino acids 1 ~ 286 did not interact with STAT3 and lead to STAT3 deactivation. In addition, MEST regulates JAK2/STAT3 complex formation through the interaction of MEST and STAT3. Moreover, high MEST expression promotes the metastasis of breast cancer in vivo.

Overall, we first identified a new molecular and functional network present in cancer metastasis that regulates and coordinates MEST. These results suggest that MEST seems to represent a new potential target for improving the treatment efficacy of metastatic cancer and the development of novel anticancer therapeutics.

## Methods

### Cell Lines and Cell Culture

Mouse (67NR, 4T07, and 4T1) and human breast cancer cells (MCF10A, MCF7, T47D, BT474, ZR75B, MDA-MB435, MDA-MB-468, SK-BR-3, BT549, Hs578T, and MDA-MB-231) and 293T cells obtained from American Type Culture Collection (Manassas, VA, USA); SUM149PT and SUM159PTPT breast cells were obtained from Dr. Stephen Ethier (Kramanos Institute) and were maintained as described previously (Fillmore & Kuperwasser, 2008; Yang et al, 2004). HMLE were obtained from Dr. Robert A. Weinberg (Massachusetts Institute of Technology) and maintained as described previously (Scheel et al, 2011). Human IL-6 were purchased from PeproTech.

### Human breast tumor samples

RNA from normal and tumorous human breast samples was obtained from the Gangnam Severance Hospital after approval (IRB approval number: 3-2011-0191) as previously described (Kim et al, 2015).

### Plasmids and viral production

pLKO.1 lentiviral plasmids encoding shRNAs targeting the MEST gene were obtained from Sigma-Aldrich (SHCLNG-NM_002402 and SHCLNG-NM_008590). Hs578T, SUM 159, and 4T1 cells were selected for 15 days with 1 mg/ml puromycin after infection with lentivirus as previously described (Kim et al, 2015). The cDNA encoding human MEST was subcloned into pEF6/V5-His-TOPO and plenti6.3/V5-TOPO vectors, respectively (Invitrogen). The cDNA encoding MESTΔC was produced via amplification of residues 1-286 of MEST using primers (**Table S1**). STAT3 deletion constructs(Kwon et al, 2008) were provided by (Young-Yun, Kong, Korea Advanced Institute of Science and Technology, Taejon, Korea).

### Antibodies, Western blotting, immunoprecipitation, and immunofluorescence

We performed Western blotting, immunoprecipitation, and immunofluorescence analysis as previously described (Jin et al, 2010). Antibodies were obtained from the following sources: Anti-MEST (SAB2501254) and anti-Flag (F3165) were from Sigma-Aldrich; anti-V5 (R960-CUS) was from Invitrogen; anti-JAK2 (ab108596), anti-Twist-1 (ab50887), anti-phosphoserine/threonine (ab17464), and anti-phosphotyrosine (ab10321) were from Abcam; anti-STAT3 (9139), anti-phospho-STAT3 (4113), and anti-phospho-JAK2 (3771) were from Cell Signaling Technology; and anti-E-cadherin (610405), anti-fibronectin (610078), anti-N-cadherin (610920), anti-alpha-catenin (610194), anti-vimentin (550513) and anti-beta-catenin (610154) were from BD Phamingen™.

### Luciferase reporter assay

Cells that were 50% confluent in 12-well dishes were transfected using Lipofectamine 2000 (Invitrogen). A total of 0.5 μg STAT3 or Twist-1 reporter gene constructs and 0.5 μg of pCMV-β-gal were co-transfected per well. The cell extracts were prepared 48 hrs after transfection, and the luciferase activity was quantified using the Enhanced Luciferase Assay Kit (BD Biosciences). All experiments were performed in triplicate.

### Soft agar assays, wound healing assays, anchorage-independent cell growth, RT-PCR, and Matrigel invasion assays

All the assays were performed as previously described (Jin et al, 2010; Kim et al, 2015). For anchorage-independent cell growth and soft agar assays, 1 × 10^3^ cells were seeded into 6 well cell culture plates. For the wound healing assay, 1 × 10^6^ cells were seeded into 6 well cell culture plates. For the invasion assay, 1 × 10^4^ cells were seeded into BD Matrigel invasion chambers with 8 μm pores (Corning, 62405-744). Additionally, the primer sequences used to amplify the genes are listed in the supplemental experimental procedures (**Table S1**).

### RNA preparation and Quantitative RT-PCR

Total RNA was isolated using RNeasy Mini Kits (Qiagen) according to the manufacturer’s instructions and reverse transcribed with hexa-nucleotide mix (Roche). The resulting cDNA was subjected to PCR using the SYBR-Green Master PCR mix and the Taqman master PCR mix (Applied Biosystems) in triplicate. PCR and data collection were performed using the 7900HT Fast Real-Time PCR System (Applied Biosystems). All quantitations were normalized to the 18S RNA endogenous control. Specific Human MEST (Hs00853380_g1), Mouse MEST (Mm00485003_m1), mouse 18S (Mm04277571_S1), and human 18S (Hs99999901_s1) quantitative probes for Taqman RT-PCR were obtained from Applied Biosystems.

### Animal studies

Animal studies were performed as previously described (Kim et al, 2015). For extravasation studies, 4T1 control-shRNA or MEST-shRNA cells (1 × 10^6^ cells) were injected into the tail vein of female BALB/c mice (7 weeks old, *n* = 7) and handled in compliance with protocols approved by the Institutional Animal Care and Use Committee (IACUC) of Gachon University (Approval No. LCDI-2010-0074). After 29 days, the lungs were excised, fixed in 10% formalin, paraffin embedded and sectioned for analysis.

### Microarray data analysis

MEST expression signatures in 2,136 breast cancer patients of the METABRIC dataset (Curtis et al, 2012) and 855 breast cancer patients of the UNC dataset (GSE26338) (Harrell et al, 2012) were extracted and averaged, and then *in silico* analysis was performed. The boxplot graphs and Kaplan-Meier survival analysis were plotted with gene expression using GraphPad Prism v 5.0 (GraphPad Software, Inc.). *P* < 0.0001 was considered statistically significant.

### Statistical analysis

Data are expressed as the means ± SEM. Statistical analyses of the data were conducted via the Student’s t test (two-tailed) and ANOVA. Differences were considered statistically significant at *P* < 0.05, *P* < 0.001, or *P* < 0.0001.

## Acknowledgements

This work was supported by a National Research Foundation of Korea grant (NRF-2010-0002525, NRF-2012R1A2A2A01002728 to W. J. and 2015R1D1A1A01059406 to MS, K). This research was a part of the project entitled ‘Development of Biomedical materials based on marine proteins’, funded by the Ministry of Oceans and Fisheries, Korea.

## Author Contributions

M.S.K., S.K.K., and W.J. designed research; M.S.K., H.S.L, Y.J.K., and D.Y.L. performed research; M.S.K., S.K.K., and W.J. analyzed data; and S.K.K and W.J. wrote the main manuscript text.

---

**Table S1.**
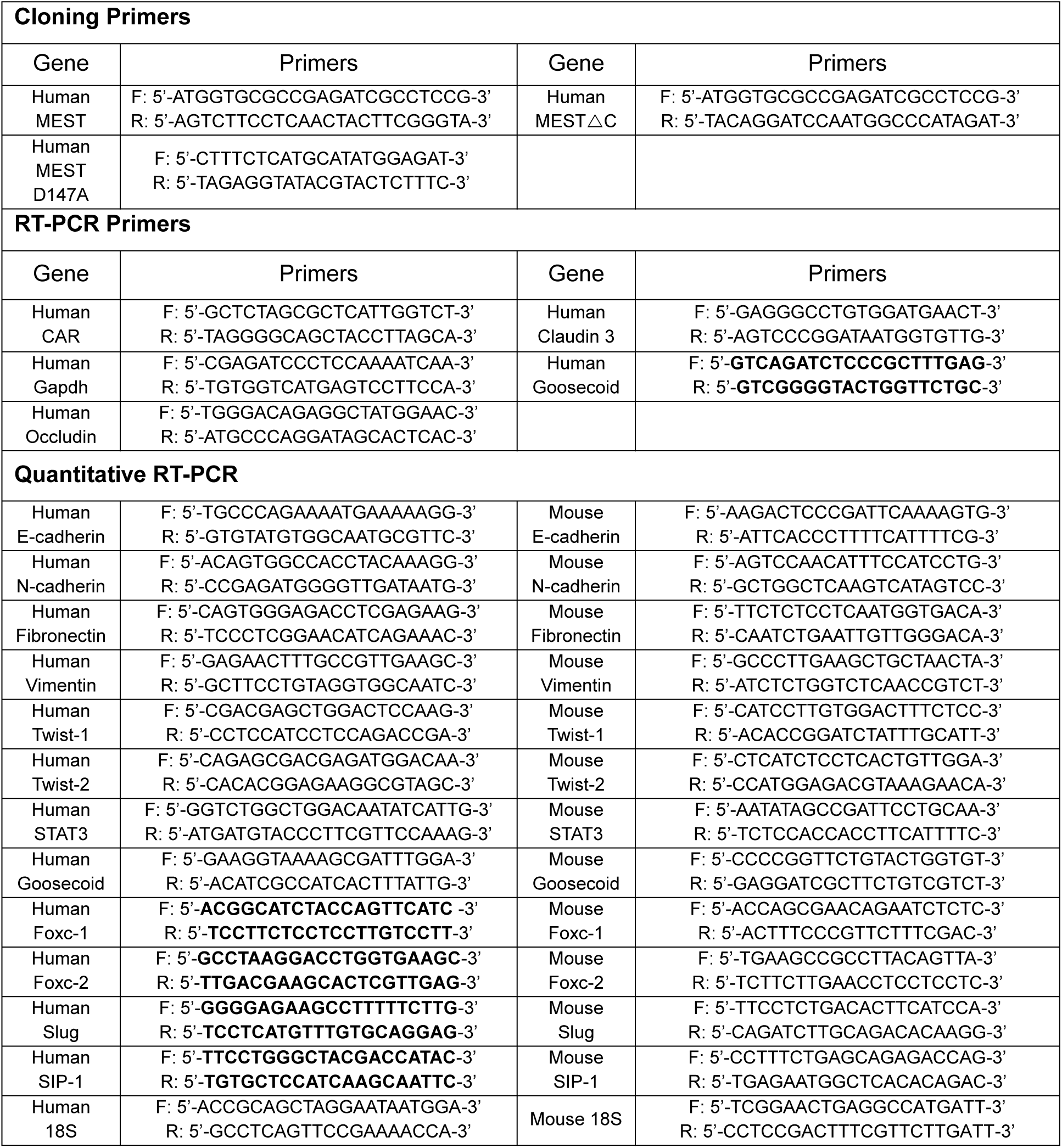
Primer sequences for Cloning, RT-PCR and quantitative RT-PCR

**Figure S1.**
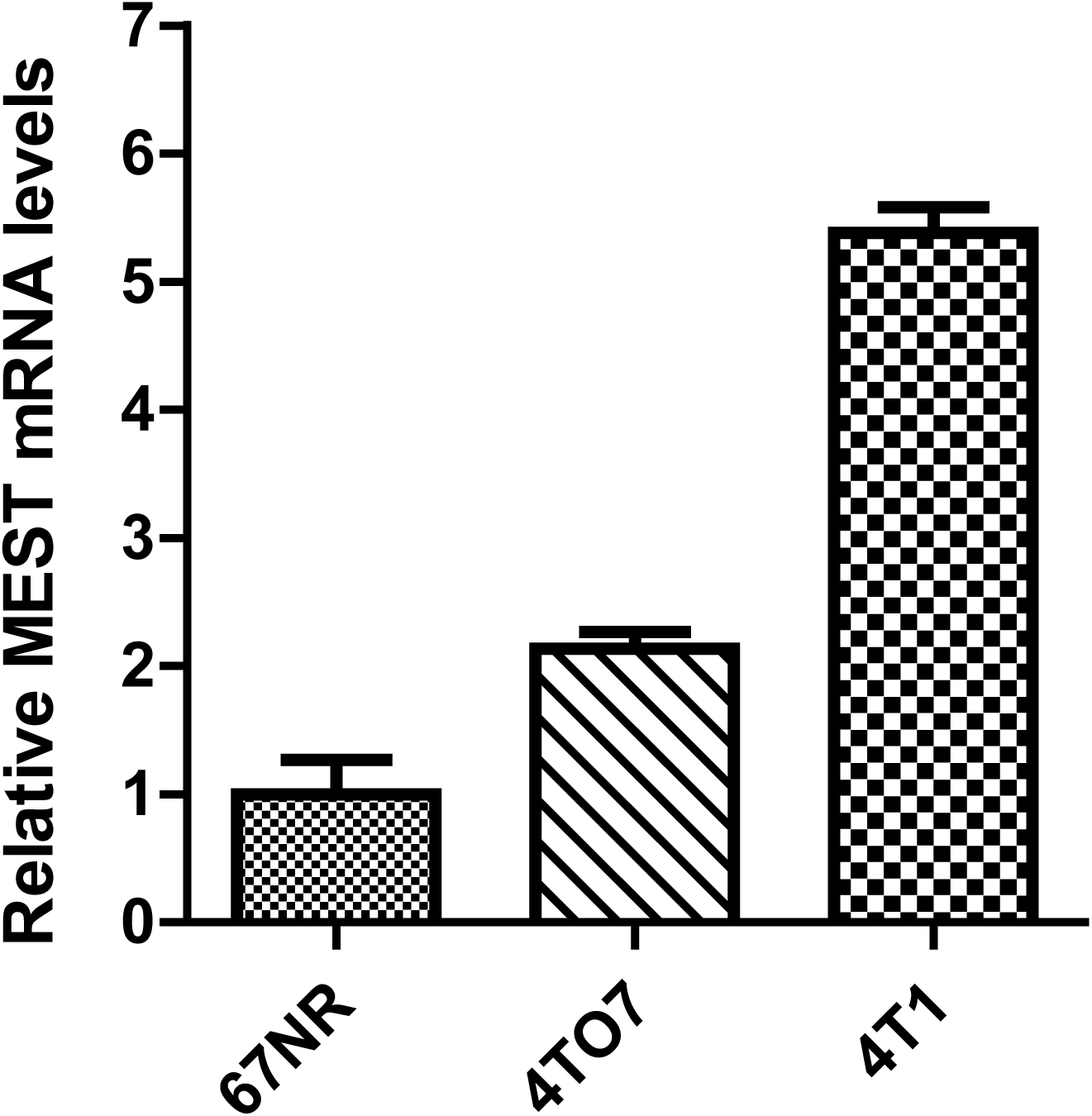
Identification of MEST expression in mouse breast cancer cells. Quantitative RT-PCR analysis of the expression of MEST in 67NR, 4T07, and 4T1 cells. GAPDH was used as a loading control. *P* < 0.001, ANOVA.

**Figure S2.**
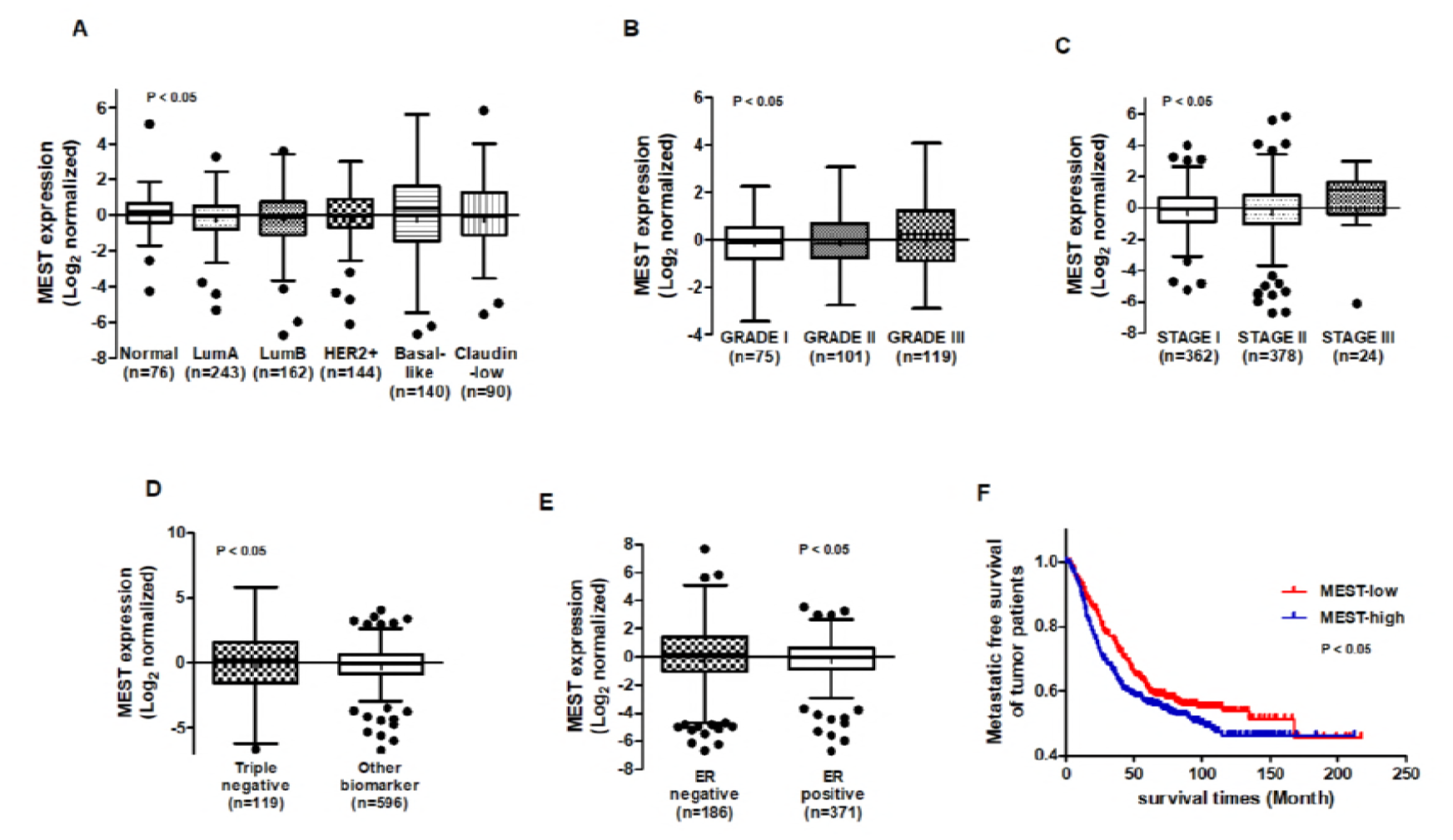
Expression of MEST is correlated with pathology of breast cancer patients. (A) Box-and-whisker (Tukey) plots are shown for the expression of MEST in 885 human breast cancer patients by breast cancer subtypes. *P* < 0.05, ANOVA. (B) Box-and-whisker (Tukey) plots are shown for the expression of MEST in the grade of 885 human breast cancer patients. *P* < 0.05, ANOVA. (C) Box-and-whisker (Tukey) plots are shown for the expression of MEST in the stage of 885 human breast cancer patients. *P* < 0.05, ANOVA. (D) Box-and-whisker (Tukey) plots are shown for the expression of MEST in triple negative or other biomarkers of 885 human breast cancer patients. *P* < 0.05, *t*-test. (E) Box-and-whisker (Tukey) plots are shown for the expression of MEST in ERα negative or ERα positive samples of the 885 human breast cancer patients. The MEST level from Figure S1A, S1B, S1C, S1D, and S1E was extracted from the UNC dataset (GSE26338) and averaged for each tumor. Points below and above the whiskers are drawn as individual dots. *P* < 0.05, *t*-test. (I) Kaplan-Meier survival plot of low and high MEST expression on UNC dataset (GSE26338). Patients were divided into high- and low-MEST expressers, and their survivals were compared. The *P* value was calculated by a log-rank test (*P* < 0.05).

**Figure S3.**
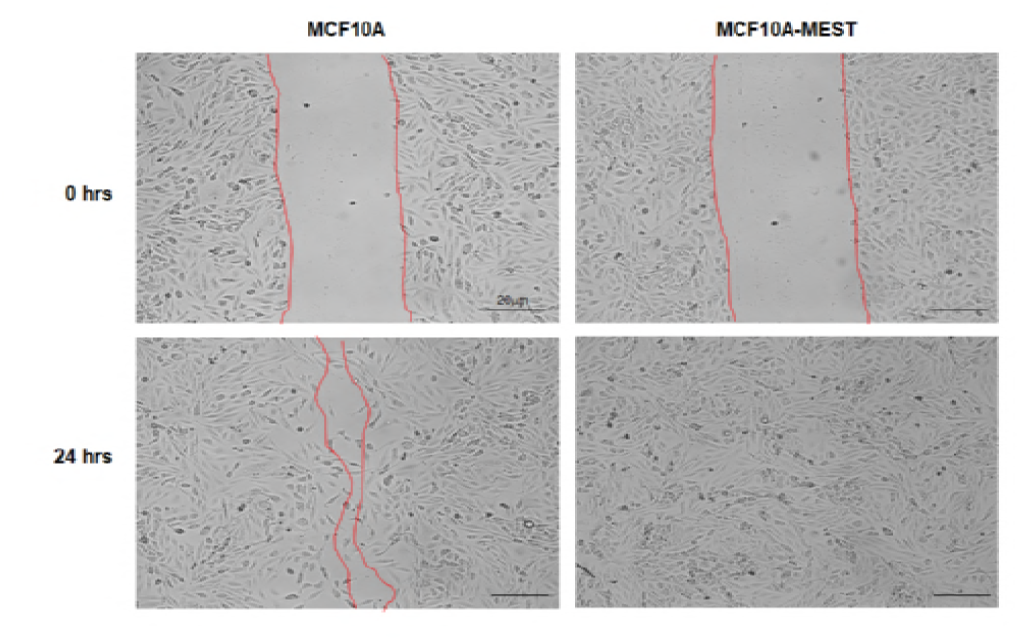
Ectopic MEST expression increases the motility of non-tumorigenic and non-metastatic MCF10A cells. Wound healing assay of the MCF10A control or MCF10A-MEST cells. Wound closures were photographed at 0 and 24 hrs after wounding.

**Figure S4.**
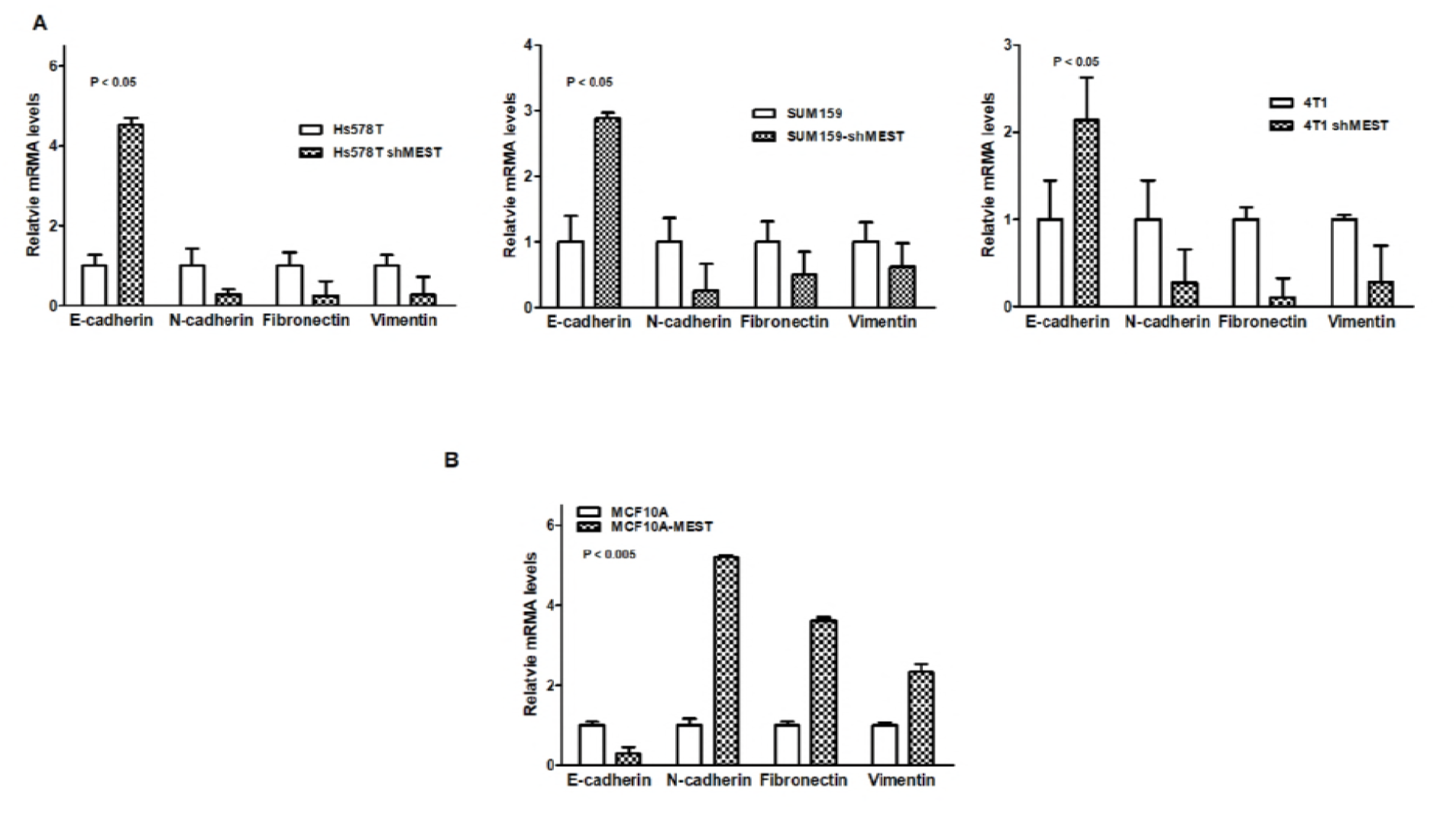
MEST regulates EMT markers. (A) The relative expression levels of mRNA encoding E-cadherin, N-cadherin, Fibronectin, and Vimentin in Hs578T, SUM159PT, and 4T1 control-shRNA or MEST-shRNA cells, as determined by quantitative RT-PCR. 18S was used as a loading control. *P* < 0.05, *t*-test. (B) The relative expression levels of mRNA encoding E-cadherin, N-cadherin, Fibronectin, and Vimentin in MCF10A control or MCF10A-MEST cells, as determined by quantitative RT-PCR. 18S was used as a loading control *P* < 0.005, *t*-test.

**Figure S5.**
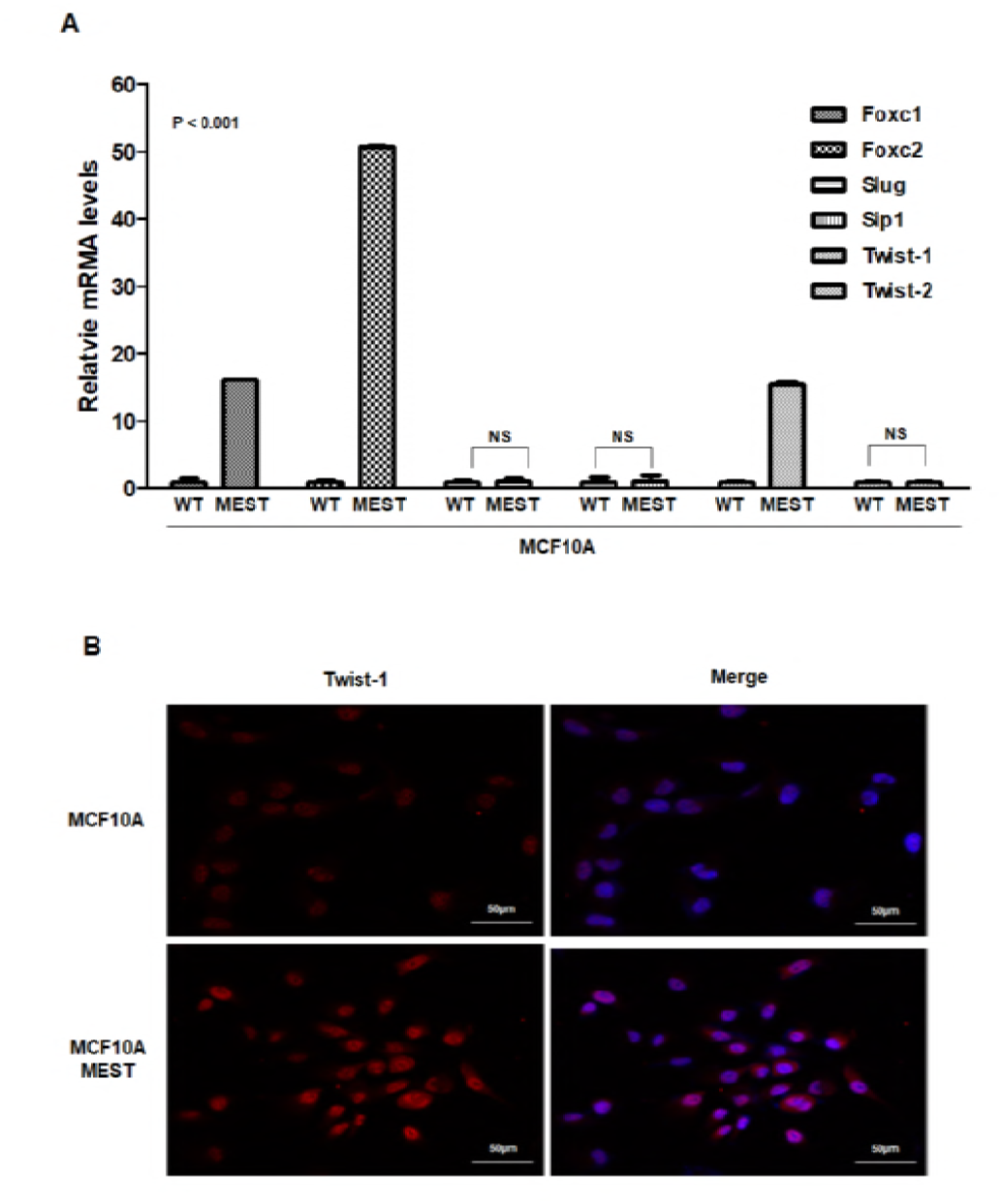
Ectopic MEST expression increases EMT-inducing transcription factors. (A) The relative expression levels of mRNA encoding Foxc-1, Foxc-2, Slug, SIP-1, Twist-1, and Twist-2 in MCF10A control or MCF10A-MEST cells, as determined by quantitative RT-PCR. 18S was used as a loading control. *P* < 0.001, NS: not significant, *t*-test. (B) Immunofluorescence images of Twist-1 in MCF10A control or MCF10A-MEST cells. The red signal represents staining of the corresponding protein, while the blue signal represents DAPI staining.

**Figure S6.**
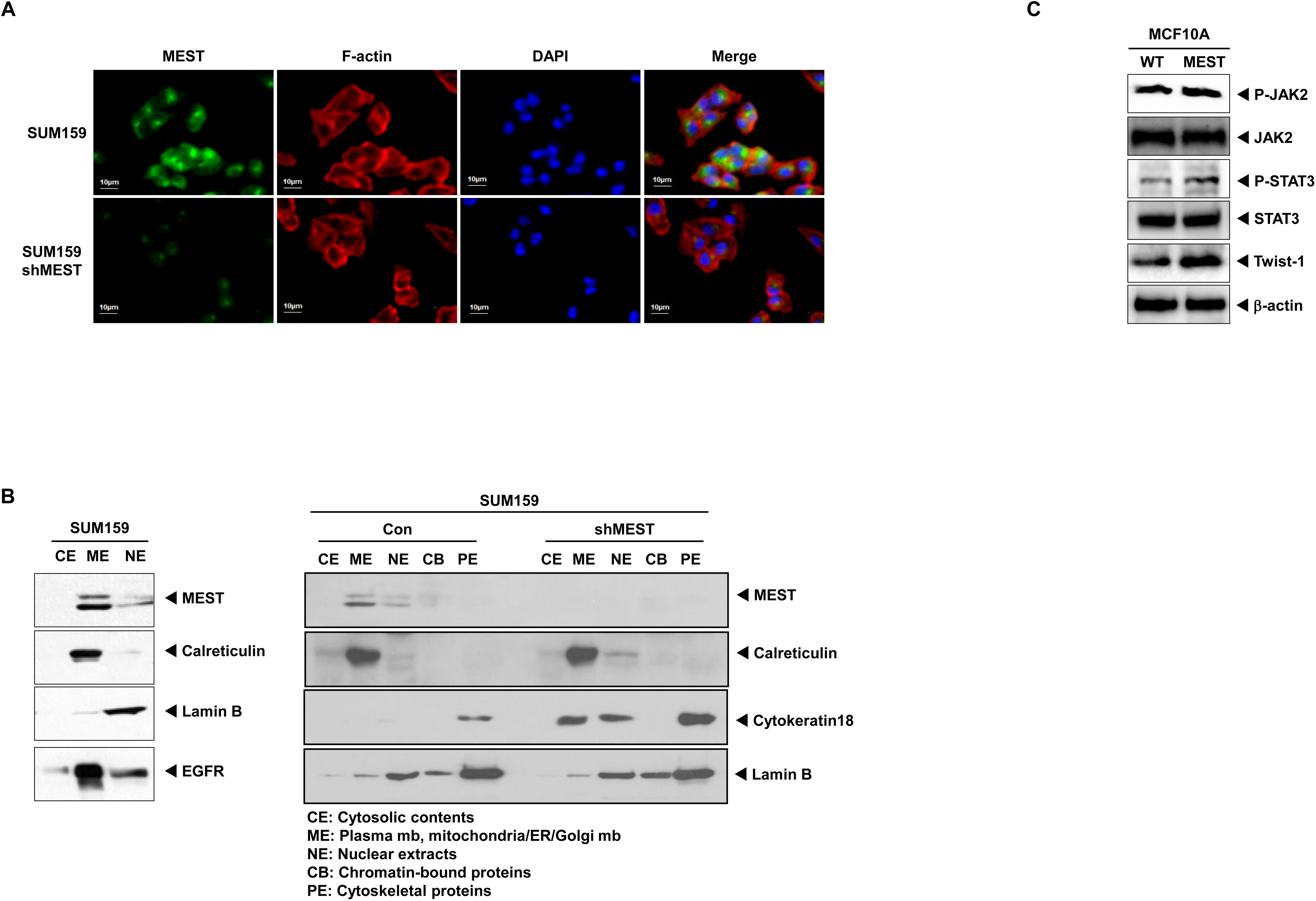
Identification of subcellular localization of MEST protein and ectopic MEST expression increases STAT3 activation and Twist-1 expression. (A) Immunofluorescence images of MEST protein in SUM159PT control-shRNA or MEST-shRNA cells. The green signal represents staining of the corresponding protein, while the blue signal represents DAPI staining and the red signal represents F-actin. (B) Western blot analysis of fractionated cellular localization of the MEST protein. Various cellular compartments, including the endoplasmic reticulum (calreticulin), nuclear soluble fraction (Lamin B), membrane/cytoplasm (EGFR) and cytoskeleton (cytokeratin 18) were used as loading controls. (C) Western blot analysis of the expression of phospho-JAK2, JAK2, phospho-STAT3, STAT3, and Twist-1 proteins in the MCF10A control or MCF10A-MEST cells. β-actin was used as a loading control.

**Figure S7.**
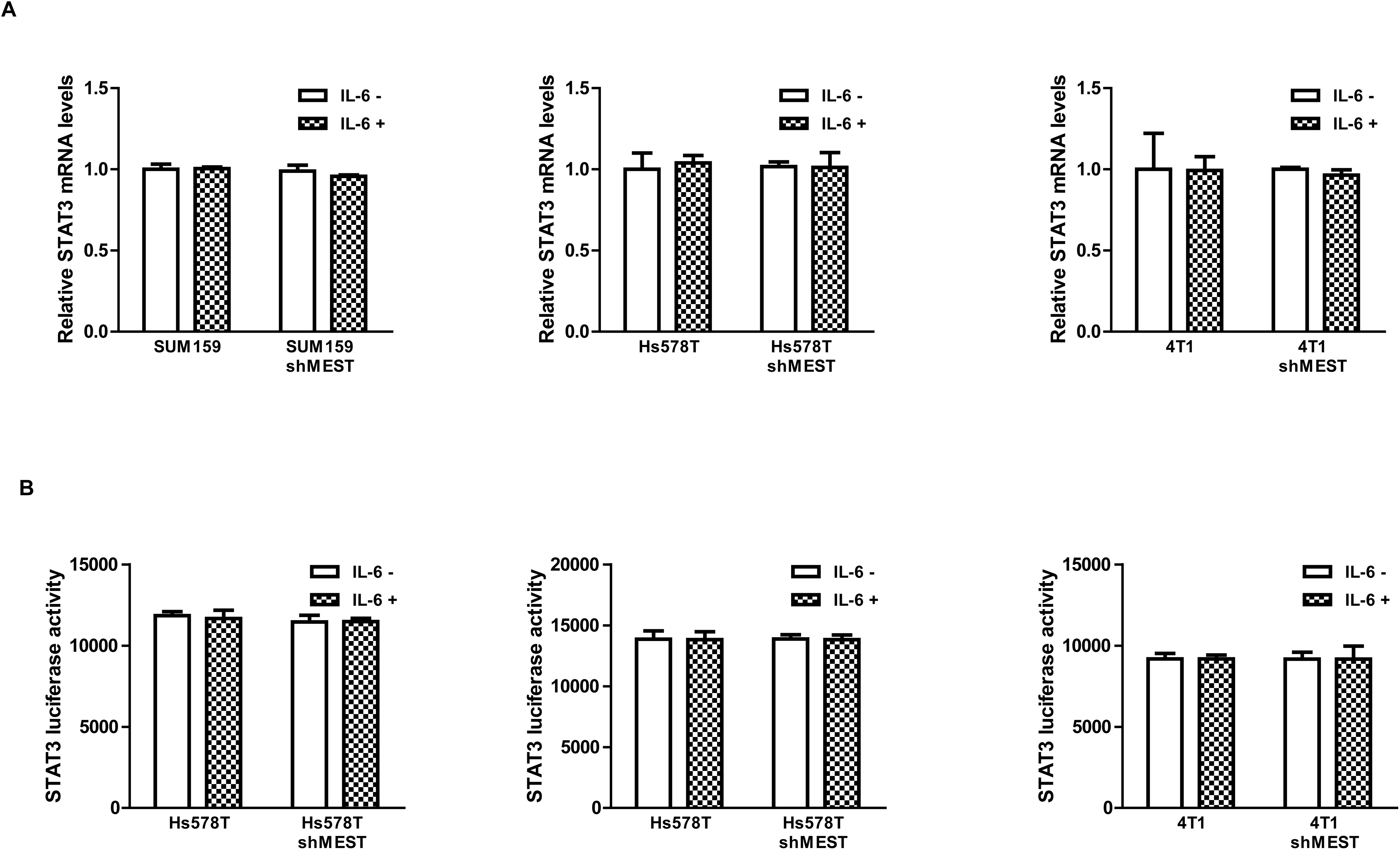
MEST expression does not affect in STAT3 mRNA levels. (A) The relative expression levels of mRNA encoding STAT3 in Hs578T, SUM159PT, and 4T1 control-shRNA or MEST-shRNA cells after treatment of IL-6 (20 ng/ml), as determined by quantitative RT-PCR. 18S was used as a loading control. NS: not significant, *t*-test. (B) Promoter activity of the STAT3 gene in Hs578T, SUM159PT, and 4T1 control-shRNA or MEST-shRNA cells after IL-6 treatment (20 ng/ml) measured by luciferase reporter assay. Each bar represents the mean ± SEM of three experiments. NS: not significant, *t*-test.

**Figure S8.**
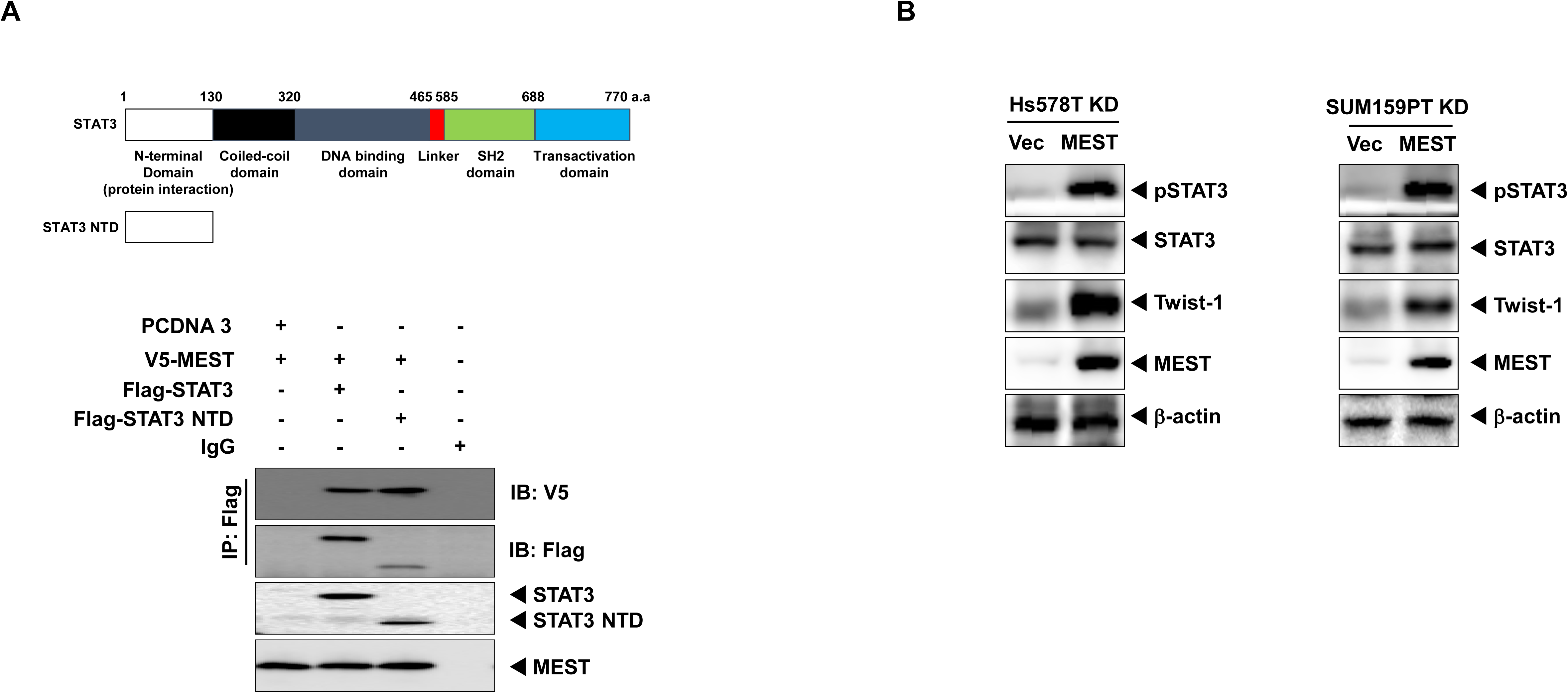
Identification of MEST region interacted with STAT3 and MEST induces STAT3 activation. (A) Western blot analysis of whole-cell lysates and immunoprecipitates derived from 293T cells transfected with the Flag-STAT3, Flag-STAT3 NTD or V5-MEST constructs, as indicated. (B) Western blot analysis of the expression of phospho-STAT3, STAT3, and Twist-1 in Hs578T, SUM159PT MEST-shRNA cells after transient transfection of MEST constructs. β-actin was used as a loading control.

## References

Ansieau S, Bastid J, Doreau A, Morel AP, Bouchet BP, Thomas C, Fauvet F, Puisieux I, Doglioni C, Piccinin S, Maestro R, Voeltzel T, Selmi A, Valsesia-Wittmann S, Caron de Fromentel C, Puisieux A (2008) Induction of EMT by twist proteins as a collateral effect of tumor-promoting inactivation of premature senescence. Cancer Cell 14: 79–89

Buettner R, Mora LB, Jove R (2002) Activated STAT signaling in human tumors provides novel molecular targets for therapeutic intervention. Clin Cancer Res 8: 945–954

Cheng GZ, Zhang WZ, Sun M, Wang Q, Coppola D, Mansour M, Xu LM, Costanzo C, Cheng JQ, Wang LH (2008) Twist is transcriptionally induced by activation of STAT3 and mediates STAT3 oncogenic function. J Biol Chem 283: 14665–14673

Creighton CJ, Li X, Landis M, Dixon JM, Neumeister VM, Sjolund A, Rimm DL, Wong H, Rodriguez A, Herschkowitz JI, Fan C, Zhang X, He X, Pavlick A, Gutierrez MC, Renshaw L, Larionov AA, Faratian D, Hilsenbeck SG, Perou CM, Lewis MT, Rosen JM, Chang JC (2009) Residual breast cancers after conventional therapy display mesenchymal as well as tumor-initiating features. Proceedings of the National Academy of Sciences of the United States of America 106: 13820–13825

Curtis C, Shah SP, Chin SF, Turashvili G, Rueda OM, Dunning MJ, Speed D, Lynch AG, Samarajiwa S, Yuan Y, Graf S, Ha G, Haffari G, Bashashati A, Russell R, McKinney S, Group M, Langerod A, Green A, Provenzano E, Wishart G, Pinder S, Watson P, Markowetz F, Murphy L, Ellis I, Purushotham A, Borresen-Dale AL, Brenton JD, Tavare S, Caldas C, Aparicio S (2012) The genomic and transcriptomic architecture of 2,000 breast tumours reveals novel subgroups. Nature 486: 346–352

Fidler IJ (2003) The pathogenesis of cancer metastasis: the ‘seed and soil’ hypothesis revisited. Nature reviews Cancer 3: 453–458

Fillmore CM, Kuperwasser C (2008) Human breast cancer cell lines contain stem-like cells that self-renew, give rise to phenotypically diverse progeny and survive chemotherapy. Breast cancer research: BCR 10: R25

Frank NY, Schatton T, Frank MH (2010) The therapeutic promise of the cancer stem cell concept. J Clin Invest 120: 41–50

Hanahan D, Weinberg RA (2011) Hallmarks of cancer: the next generation. Cell 144: 646–674

Harrell JC, Prat A, Parker JS, Fan C, He X, Carey L, Anders C, Ewend M, Perou CM (2012) Genomic analysis identifies unique signatures predictive of brain, lung, and liver relapse. Breast Cancer Res Treat 132: 523–535

Hartwell KA, Muir B, Reinhardt F, Carpenter AE, Sgroi DC, Weinberg RA (2006) The Spemann organizer gene, Goosecoid, promotes tumor metastasis. Proceedings of the National Academy of Sciences of the United States of America 103: 18969–18974

Haura EB, Turkson J, Jove R (2005) Mechanisms of disease: Insights into the emerging role of signal transducers and activators of transcription in cancer. Nat Clin Pract Oncol 2: 315–324

Hennessy BT, Gonzalez-Angulo AM, Stemke-Hale K, Gilcrease MZ, Krishnamurthy S, Lee JS, Fridlyand J, Sahin A, Agarwal R, Joy C, Liu W, Stivers D, Baggerly K, Carey M, Lluch A, Monteagudo C, He X, Weigman V, Fan C, Palazzo J, Hortobagyi GN, Nolden LK, Wang NJ, Valero V, Gray JW, Perou CM, Mills GB (2009) Characterization of a naturally occurring breast cancer subset enriched in epithelial-to-mesenchymal transition and stem cell characteristics. Cancer research 69: 4116–4124

Jin W, Kim GM, Kim MS, Lim MH, Yun C, Jeong J, Nam JS, Kim SJ (2010) TrkC plays an essential role in breast tumor growth and metastasis. Carcinogenesis 31: 1939–1947

Kim MS, Kim GM, Choi YJ, Kim HJ, Kim YJ, Jin W (2013a) c-Src activation through a TrkA and c-Src interaction is essential for cell proliferation and hematological malignancies. Biochem Biophys Res Commun 441: 431–437

Kim MS, Kim GM, Choi YJ, Kim HJ, Kim YJ, Jin W (2013b) TrkC promotes survival and growth of leukemia cells through Akt-mTOR-dependent up-regulation of PLK-1 and Twist-1. Mol Cells 36: 177–184

Kim MS, Lee WS, Jeong J, Kim SJ, Jin W (2015) Induction of metastatic potential by TrkB via activation of IL6/JAK2/STAT3 and PI3K/AKT signaling in breast cancer. Oncotarget 6: 40158–40171

Kosaki K, Kosaki R, Craigen WJ, Matsuo N (2000) Isoform-specific imprinting of the human PEG1/MEST gene. Am J Hum Genet 66: 309–312

Kwok WK, Ling MT, Lee TW, Lau TC, Zhou C, Zhang X, Chua CW, Chan KW, Chan FL, Glackin C, Wong YC, Wang X (2005) Up-regulation of TWIST in prostate cancer and its implication as a therapeutic target. Cancer research 65: 5153–5162

Kwon MC, Koo BK, Moon JS, Kim YY, Park KC, Kim NS, Kwon MY, Kong MP, Yoon KJ, Im SK, Ghim J, Han YM, Jang SK, Shong M, Kong YY (2008) Crif1 is a novel transcriptional coactivator of STAT3. The EMBO journal 27: 642–653

Lefebvre L, Viville S, Barton SC, Ishino F, Surani MA (1997) Genomic structure and parent-of-origin-specific methylation of Peg1. Hum Mol Genet 6: 1907–1915

Levy DE, Darnell JE, Jr. (2002) Stats: transcriptional control and biological impact. Nat Rev Mol Cell Biol 3: 651–662

Li X, Lewis MT, Huang J, Gutierrez C, Osborne CK, Wu MF, Hilsenbeck SG, Pavlick A, Zhang X, Chamness GC, Wong H, Rosen J, Chang JC (2008) Intrinsic resistance of tumorigenic breast cancer cells to chemotherapy. Journal of the National Cancer Institute 100: 672–679

Liu S, Wicha MS (2010) Targeting breast cancer stem cells. J Clin Oncol 28: 4006–4012

Ma XJ, Salunga R, Tuggle JT, Gaudet J, Enright E, McQuary P, Payette T, Pistone M, Stecker K, Zhang BM, Zhou YX, Varnholt H, Smith B, Gadd M, Chatfield E, Kessler J, Baer TM, Erlander MG, Sgroi DC (2003) Gene expression profiles of human breast cancer progression. Proceedings of the National Academy of Sciences of the United States of America 100: 5974–5979

Mani SA, Guo W, Liao MJ, Eaton EN, Ayyanan A, Zhou AY, Brooks M, Reinhard F, Zhang CC, Shipitsin M, Campbell LL, Polyak K, Brisken C, Yang J, Weinberg RA (2008) The epithelial-mesenchymal transition generates cells with properties of stem cells. Cell 133: 704–715

Mani SA, Yang J, Brooks M, Schwaninger G, Zhou A, Miura N, Kutok JL, Hartwell K, Richardson AL, Weinberg RA (2007) Mesenchyme Forkhead 1 (FOXC2) plays a key role in metastasis and is associated with aggressive basal-like breast cancers. Proceedings of the National Academy of Sciences of the United States of America 104: 10069–10074

Marotta LL, Almendro V, Marusyk A, Shipitsin M, Schemme J, Walker SR, Bloushtain-Qimron N, Kim JJ, Choudhury SA, Maruyama R, Wu Z, Gonen M, Mulvey LA, Bessarabova MO, Huh SJ, Silver SJ, Kim SY, Park SY, Lee HE, Anderson KS, Richardson AL, Nikolskaya T, Nikolsky Y, Liu XS, Root DE, Hahn WC, Frank DA, Polyak K (2011) The JAK2/STAT3 signaling pathway is required for growth of CD44(+)CD24(-) stem cell-like breast cancer cells in human tumors. The Journal of clinical investigation 121: 2723–2735

Mayer W, Hemberger M, Frank HG, Grummer R, Winterhager E, Kaufmann P, Fundele R (2000) Expression of the imprinted genes MEST/Mest in human and murine placenta suggests a role in angiogenesis. Dev Dyn 217: 1–10

Moon YS, Park SK, Kim HT, Lee TS, Kim JH, Choi YS (2010) Imprinting and expression status of isoforms 1 and 2 of PEG1/MEST gene in uterine leiomyoma. Gynecol Obstet Invest 70: 120–125

Nakanishi H, Suda T, Katoh M, Watanabe A, Igishi T, Kodani M, Matsumoto S, Nakamoto M, Shigeoka Y, Okabe T, Oshimura M, Shimizu E (2004) Loss of imprinting of PEG1/MEST in lung cancer cell lines. Oncol Rep 12: 1273–1278

Nishihara S, Hayashida T, Mitsuya K, Schulz TC, Ikeguchi M, Kaibara N, Oshimura M (2000) Multipoint imprinting analysis in sporadic colorectal cancers with and without microsatellite instability. Int J Oncol 17: 317–322

Ohuchida K, Mizumoto K, Ohhashi S, Yamaguchi H, Konomi H, Nagai E, Yamaguchi K, Tsuneyoshi M, Tanaka M (2007) Twist, a novel oncogene, is upregulated in pancreatic cancer: clinical implication of Twist expression in pancreatic juice. Int J Cancer 120: 1634–1640

Pedersen IS, Dervan PA, Broderick D, Harrison M, Miller N, Delany E, O’Shea D, Costello P, McGoldrick A, Keating G, Tobin B, Gorey T, McCann A (1999) Frequent loss of imprinting of PEG1/MEST in invasive breast cancer. Cancer Res 59: 5449–5451

Raval A, Lucas DM, Matkovic JJ, Bennett KL, Liyanarachchi S, Young DC, Rassenti L, Kipps TJ, Grever MR, Byrd JC, Plass C (2005) TWIST2 demonstrates differential methylation in immunoglobulin variable heavy chain mutated and unmutated chronic lymphocytic leukemia. J Clin Oncol 23: 3877–3885

Riesewijk AM, Hu L, Schulz U, Tariverdian G, Hoglund P, Kere J, Ropers HH, Kalscheuer VM (1997) Monoallelic expression of human PEG1/MEST is paralleled by parent-specific methylation in fetuses. Genomics 42: 236–244

Rosen JM, Jordan CT (2009) The increasing complexity of the cancer stem cell paradigm. Science 324: 1670–1673

Scheel C, Eaton EN, Li SH, Chaffer CL, Reinhardt F, Kah KJ, Bell G, Guo W, Rubin J, Richardson AL, Weinberg RA (2011) Paracrine and autocrine signals induce and maintain mesenchymal and stem cell states in the breast. Cell 145: 926–940

Taube JH, Herschkowitz JI, Komurov K, Zhou AY, Gupta S, Yang J, Hartwell K, Onder TT, Gupta PB, Evans KW, Hollier BG, Ram PT, Lander ES, Rosen JM, Weinberg RA, Mani SA (2010) Core epithelial-to-mesenchymal transition interactome gene-expression signature is associated with claudin-low and metaplastic breast cancer subtypes. Proceedings of the National Academy of Sciences of the United States of America 107: 15449–15454

Valastyan S, Weinberg RA (2011) Tumor metastasis: molecular insights and evolving paradigms. Cell 147: 275–292

Valsesia-Wittmann S, Magdeleine M, Dupasquier S, Garin E, Jallas AC, Combaret V, Krause A, Leissner P, Puisieux A (2004) Oncogenic cooperation between H-Twist and N-Myc overrides failsafe programs in cancer cells. Cancer Cell 6: 625–630

Yang J, Mani SA, Donaher JL, Ramaswamy S, Itzykson RA, Come C, Savagner P, Gitelman I, Richardson A, Weinberg RA (2004) Twist, a master regulator of morphogenesis, plays an essential role in tumor metastasis. Cell 117: 927–939

Zhang Y, Hassan MQ, Li ZY, Stein JL, Lian JB, van Wijnen AJ, Stein GS (2008) Intricate gene regulatory networks of helix-loop-helix (HLH) proteins support regulation of bone-tissue related genes during osteoblast differentiation. J Cell Biochem 105: 487–496

Zhang Z, Xie D, Li X, Wong YC, Xin D, Guan XY, Chua CW, Leung SC, Na Y, Wang X (2007) Significance of TWIST expression and its association with E-cadherin in bladder cancer. Hum Pathol 38: 598–606

